# Pangolin genomes offer key insights and resources for the world’s most trafficked wild mammals

**DOI:** 10.1101/2023.02.16.528682

**Authors:** Sean P. Heighton, Rémi Allio, Jérôme Murienne, Jordi Salmona, Hao Meng, Céline Scornavacca, Armanda D.S. Bastos, Flobert Njiokou, Darren W. Pietersen, Marie-Ka Tilak, Shu-Jin Luo, Frédéric Delsuc, Philippe Gaubert

**Affiliations:** Laboratoire Evolution et Diversité Biologique (EDB)— IRD-UPS-CNRS, Université Toulouse III, 118 route de Narbonne, Bât. 4R1, 31062 Toulouse, France; ISEM, Université de Montpellier, CNRS, IRD, Montpellier, France; School The State Key Laboratory of Protein and Plant Gene Research of Life Sciences, Peking-Tsinghua Center for Life Sciences, Peking University, Beijing 100871, China; Mammal Research Institute, Department of Zoology & Entomology, University of Pretoria, Pretoria 0002, South Africa; Laboratoire de Parasitologie et Ecologie, Université de Yaoundé I, Faculté des Sciences, Cameroon; CIIMAR/CIMAR, Centro Interdisciplinar de Investigação Marinha e Ambiental, Universidade do Porto, Terminal de Cruzeiros do Porto de Leixões, Av. General Norton de Matos, s/n, 4450-208 Porto, Portugal

**Author notes:** These authors contributed equally (Sean P. Heighton, Rémi Allio). Senior author (Shu-Jin Luo, Frédéric Delsuc, Philippe Gaubert).

**Keywords:** Pangolin, Pholidota, genomics, phylogeny, full genomes, genomic diversity, conservation, PSMC, ancient admixture, ghost lineage, divergence estimates, forensic markers, wildlife trade

## Abstract

Pangolins form a group of scaly mammals that are trafficked at record numbers for their meat and medicinal properties. Despite their great conservation concern, knowledge of their evolution is limited by a paucity of genomic data. We aim to produce exhaustive genomic resources that include 3 238 orthologous genes and whole-genome polymorphisms to assess the evolution of all eight pangolin species. Robust orthologous gene-based phylogenies recovered the monophyly of the three genera of pangolins, and highlighted the existence of an undescribed species closely related to South-East Asian pangolins. Signatures of middle Miocene admixture between an extinct, possibly European, lineage and the ancestor of South-East Asian pangolins, provides new insights into the early evolutionary history of the group. Demographic trajectories and genome-wide heterozygosity estimates revealed contrasts between continental *vs*. island populations and species lineages, suggesting that conservation planning should consider intra-specific patterns. With the expected loss of genomic diversity from recent, extensive trafficking not yet been realized in pangolins, we recommend that populations are genetically surveyed to anticipate any deleterious impact of the illegal trade. Finally, we produce a complete set of genomic resources that will be integral for future conservation management and forensic endeavors required for conserving pangolins, including tracing their illegal trade. These include the completion of whole-genomes for pangolins through the first reference genome with long reads for the giant pangolin (*Smutsia gigantea*) and new draft genomes (~43x–77x) for four additional species, as well as a database of orthologous genes with over 3.4 million polymorphic sites.

## Introduction

Genomics is becoming a standard in wildlife research as it provides genome-wide data for more accurate inferences on species or population delimitation, demographic parameters, diversity, historical trajectories, and the adaptive capacity to global changes (Allendorf, et al. 2010). Although transforming this research into conservation practice is yet to be common (Shafer, et al. 2015), the gap is closing (Garner, et al. 2016; Formenti, et al. 2022; Paez, et al. 2022).

Pangolins are a group of mammals harboring eight extant species (four in Africa and Asia) under the order Pholidota (Gaubert, et al. 2020) that have become a taxon of great public interest and conservation concern in recent years (Pietersen and Challender 2020; Heighton and Gaubert 2021). This is mainly due to them being the most trafficked wild mammals on Earth (Heinrich, et al. 2017) and a recent, incorrect, suggestion that they may have been linked to the COVID-19 pandemic (Frutos, et al. 2020; Lam, et al. 2020; Lee, et al. 2020). Despite their dire conservation circumstances, pangolins are considered to be understudied with major gaps in basic species or population research (Pietersen and Challender 2020; Heighton and Gaubert 2021). Even with a long-standing interest in their taxonomy, particularly in light of their convergent evolution with South American anteaters (Xenarthra: Miyamoto and Goodman 1986; Wyss, et al. 1987; Murphy, et al. 2001; Delsuc, et al. 2002), phylogenetic studies focusing on pangolins have been incomplete. They have either been restricted to single markers or mitochondrial genomes, limited by taxon coverage, or clouded by incorrect taxonomic sampling (Yu, et al. 2011; Du Toit, et al. 2014; Gaubert and Antunes 2015; Hassanin, et al. 2015; du Toit, et al. 2017). The most comprehensive phylogenetic study to date was inferred using a dataset including 15 mitogenomes and nine nuclear genes encompassing all eight species (Gaubert, et al. 2018).

As for genome-wide inferences for pangolins, the majority of studies are non-NGS (next generation sequencing) based through traditional methods like karyotyping (Ray-Chaudhuri, et al. 1969; Su, et al. 1994; Che, et al. 2008; Zhihai, et al. 2016). In fact, there are only the two Asian species (*Manis pentadactyla* and *M. javanica*) with published full genomes, both of which have become the subject of crucial conservation genomic studies (Choo, et al. 2016; Hu, Hao, et al. 2020). These studies have mainly focused on demographic history to determine the consequences of recent population fluctuations due to climatic oscillations (Hu, Hao, et al. 2020; Wei, et al. 2022). However, they have also delved into comparisons of diversity, inbreeding, mutational load amongst populations as well as population structuring for geographic assignment of illegally traded individuals (Hu, Hao, et al. 2020; Wei, et al. 2022). Despite the thoroughness of these studies on the two species, our understanding of the evolution of pangolins is still taxonomically limited, which compromises the potential of utilizing genetic markers for conservation and management purposes of the entire order (Allendorf, et al. 2010; Kotze, et al. 2020).

The genomics of pangolins is a challenging task. First, their elusive behaviour and tropical distributions render genetic sampling to be time-consuming and costly. Second, the phylogenetic isolation of the group from other orders, the limited fossil records and the deep divergence within pangolins (Gaudin, et al. 2009; Gaubert, et al. 2018), pose methodological hurdles. Sister to Pholidota is order Carnivora which are estimated to have started to diverge from each other from around 76 Million years ago (Zhou, et al. 2011; Gaubert, et al. 2018), thus making it difficult to incorporate outgroup taxa for genomic inferences of divergence and introgression. Compounded by this is the deep divergence (± 37.9 Million years ago) of Asian and African pangolins (Gaubert, et al. 2018) which may introduce ascertainment bias to genomic inferences depending on which clade the reference genome is part of for alignment of the group (Günther and Nettelblad 2019; Bohling 2020). Therefore, alternative approaches to the simple variant-based phylogenomic analyses based on the alignment of all taxa to a single pangolin reference genome need to be considered (Kapli, et al. 2020; Prasad, et al. 2022).

The basis of conservation genomics relies upon generating genomic data such as reference genomes and datasets of markers (e.g. homologous / orthologous genes, SNPs) used to make species and population inferences for application in conservation management (Allendorf, et al. 2010; Formenti, et al. 2022; Paez, et al. 2022). With pangolins in need of conservation action, compounded by limited genomic information on the group, a gap is needed to be filled. We thus aim to conduct the first full genome analysis on all eight pangolin species, to determine evolutionary relationships and demographic trends, identify key genetic parameters for species management, and develop a set of markers for future conservation genetic / genomic efforts. Additionally, we provide a high quality reference genome of the giant pangolin (*Smutsia gigantea*), an elusive fossorial species found in western and central Africa, which promises to be a key reference for the *Smutsia* genus and the African species alike.

## Results and Discussion

We sequenced and assembled the first reference genome for the genus *Smutsia* (giant pangolin; *S. gigantea*, ~87x) using long Nanopore reads and draft Illumina genomes for the black-bellied (*Phataginus tetradactyla*, ~43x), Temminck’s (*Smutsia temminckii*, ~44x), Indian (*Manis crassicaudata*, ~53x), and Philippine (*Manis culionensis*, ~77x) pangolins (Table S1). These new genomic data along with previously published reference genomes of the remaining three species, namely the white-bellied (*Phataginus tricuspis*), Sunda (*M. javanica*), and Chinese (*M. pentadactyla*) pangolins (Table S1), provide the first complete set of genomes for Pholidota. We also included a recently published genome of a pangolin of uncertain origin and taxonomy, seized from south-western China (Cao, et al. 2021). In an effort to reduce ascertainment bias from mapping deeply divergent Asian and African species to a single reference (Albrechtsen, et al. 2010; Lachance and Tishkoff 2013; Prasad, et al. 2022), species were mapped to a representative genome from their respective clade (Figure S1). The mapped genomes then underwent haploid and IUPAC ambiguity codes (diploid) consensus assignment before using mtDNA haplotypes and 3 239 entire single-copy orthologous autosomal genes (introns and exons) for phylogenomic and divergence time inferences.

Our partitioned concatenated (supermatrix), non-partitioned concatenated, and coalescent (summary tree) phylogenies based on orthologous, whole-gene markers show robust support for the previously reported clear dichotomy between African and Asian pangolins, and the three main clades in agreement with the three distinct pangolin genera (Figures 1 & S2; Gaubert, et al. 2018). The seized individual from Sichuan China, either identified as *M. culionensis* in Cao, et al. (2021) –based on mitogenomic data– or *M. crassicaudata* on NCBI (GCA_016801295), is sister to the South-East Asian pangolins (*M. javanica* and *M. culionensis*) and does not branch with the Indian pangolin (*M. crassicaudata*). Our mitochondrial phylogenies support the orthologous full nuclear gene phylogeny (Figure S3a), while the *Cytb* (Figure S3b) and *COI* (Figure S3c) gene phylogenies suggest that this individual is forming a monophyletic clade with two samples seized in Hong Kong. These two samples, with which this individual nests, have been suggested to be a potentially new Asian pangolin species based on *Cytb* and *COI* species delimitation (Zhang, et al. 2015; Hu, Roos, et al. 2020). Our results using orthologous whole-gene phylogenies with high bootstrap support and concordance analysis provide additional evidence that this individual likely represents a separate Asian species. However, with high levels of cryptic diversity across pangolin species (Gaubert, et al. 2016; Gaubert, et al. 2018), particularly within *M. culionensis* and *M. javanica* (Zhang, et al. 2015; Nash, et al. 2018; Hu, Roos, et al. 2020), we cannot be certain that this new taxon is indeed a new *Manis* species or part of the *M. javanica* species complex (including *M. culionensis*). Additional analyses using geo-referenced wild samples, as well as species testing through genomic and morphological data will provide further insight into the status of this taxon and its distribution. These parameters will be crucial in determining conservation priorities and management plans for this potentially new species.

**Figure 1:**
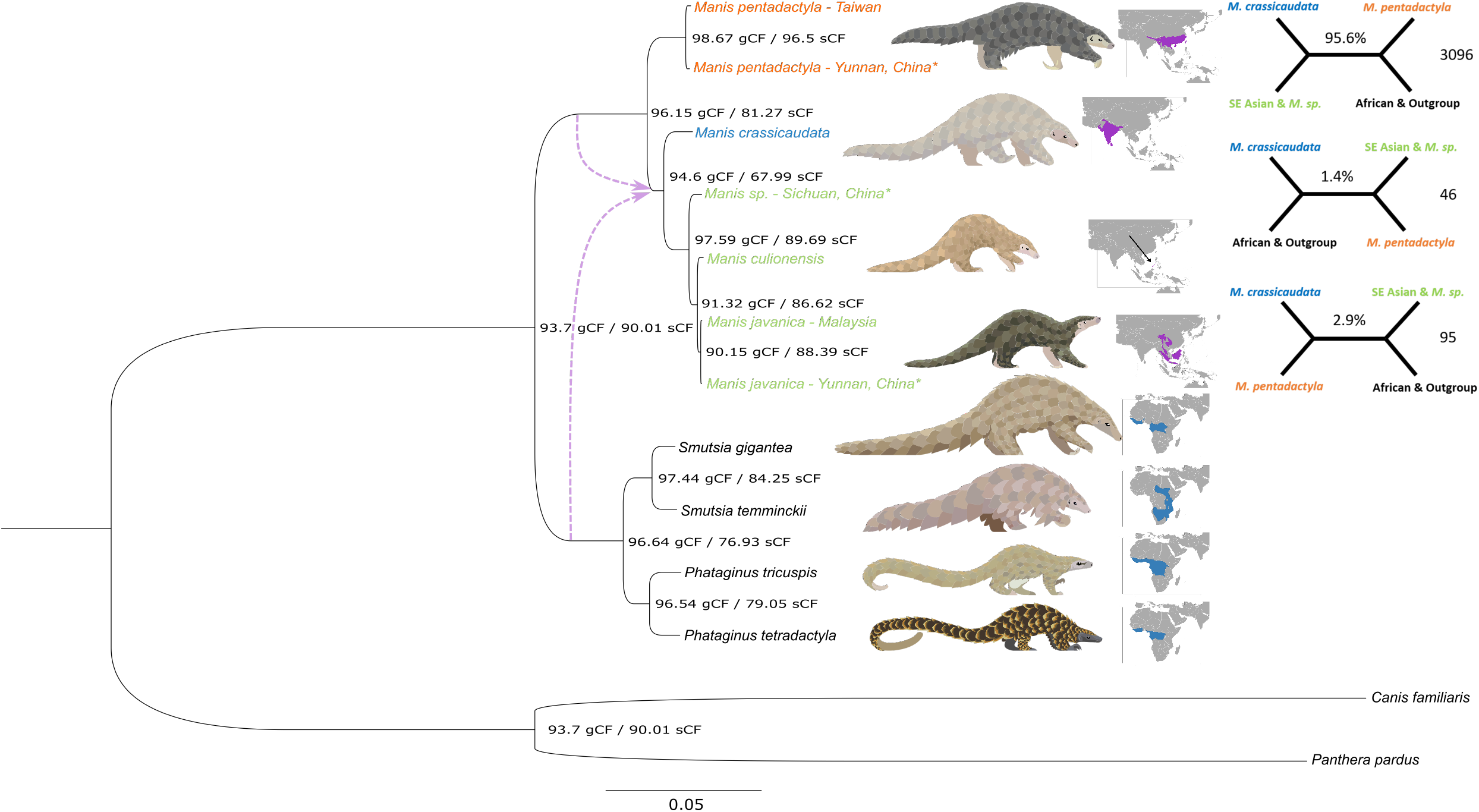
Phylogenomic relationships and reticulation amongst pangolins. Phylogenetic tree based on the alignment(s) of 58 724 014 bp from 2 238 IUPAC consensus whole-gene markers. The pangolin phylogeny consists of 13 individuals from all eight species and is rooted with two representatives of the sister order Carnivora (*Canis familiaris* and *Panthera pardus*). The same topology was derived from a partitioned concatenated (supermatrix), non-partitioned concatenated (Figure S2a) and coalescent (summary tree; Figure S2b) phylogenies, hence only the former is shown. All three phylogenies were derived from an orthologous whole-gene pipeline (Figure S1) and had full branch support for all nodes (100 for the concatenated phylogenies using 1 000 Felsenstein bootstrap replicates and 1 for the coalescent phylogeny using local posterior probability). Nodal numbers represent the concordance factors of genes (gCF) and sites (sCF), which indicate the proportion of genes and sites that fit the current topology. sCF values were based on 100 quartets randomly sampled around each internal branch. Dotted lines (purple) indicate a suggested reticulation event either from African or Asian basal nodes based on maximum pseudolikelihood phylonetworks testing (Figure S4). The main species tree quartet and the two alternative gene tree quartet topologies (from top to bottom on the top right) were identified from the coalescent analysis (Figure S2c). These quartets only implicate the internal branch around the four Asian pangolin species in corresponding colors in the main phylogeny. The value at the center of each quartet refers to the proportion of gene trees following this quartet while the value below the branch is the corresponding number of gene trees. Asterisks (*) indicate confiscated individuals whose origins could not be verified. Pangolin illustrations by Sheila McCabe.

### Reticulation and incomplete lineage sorting in South-East Asian pangolins

To assess the level of alternative topologies across the phylogenetic species tree, we used a gene and site concordance analysis, along with the proportion of alternative quartets around each branch of the main topology as obtained from the coalescent species tree analysis (Zhang, et al. 2018; Minh, Hahn, et al. 2020). Overall, there was a low level of discordance and proportion of alternative topologies (d = 0.981; Figures 1 & S2c), indicating a global robustness of the pangolin tree topology (Minh, Hahn, et al. 2020). The clade containing *M. javanica* and *M. culionensis* had the highest values for both gene discordance (90.6 gCF / 88.4 sCF) and alternative quartet topologies (8.81%; Figures 1 & S2c). *Manis javanica* and *M. culionensis* are currently regarded as distinct species based on five discriminant morphological characters (Gaubert and Antunes 2005) and phylogenetic species delimitation (Gaubert, et al. 2018), although mean pairwise genetic distances between the two were lower than any other species combinations, including estimates across the six *P. tricuspis* lineages. Thus, our results demonstrate the need for extensive population-level estimates across the distributions of South-East Asian pangolin species, including the new *Manis sp*. taxon, to aid in species conservation planning and accurate post-seizure repatriation of live individuals (Nash, et al. 2018). The latter is particularly pertinent given the multiple South-East Asian islands from which lineages of *M. javanica* are sourced and traded (Zhang, et al. 2015; Nash, et al. 2018).

Given the relatively short branch length possibly causing this uncertainty in separating *M. culionensis* from *M. javanica* through incomplete lineage sorting (ILS; Blom, et al. 2016), we conducted a chi-squared test on the gene concordance factors. The results suggest that ILS is the sole cause of discordance as the alternative topologies are not significantly independent (p <0.05; Table S2) regarding the frequency of gene trees supporting each topology (Huson, et al. 2005; Zheng and Janke 2018). This is however not the case for the branch leading to the *M. crassicaudata*/*M. sp*./*M. culionensis*/*M. javanica* clade whereby alternative topologies are significantly independent (Figures 1 & Table S2). The clade also is an outlier for the lowest site concordance (94.6 gCF / 67.99 sCF), which points to site discrepancies for this clade. With introgression (ancient gene flow) and hybridization being a possible cause of unbalanced gene-tree discordance of this clade (Doyle 1992), we tested for this using a maximum pseudolikelihood network reticulation analysis across the entire topology (Than, et al. 2008). This was conducted on all 3 238 gene trees. The results indicate that no more than one reticulation event is the most likely for pangolins (Figure 1), and that the timing and direction of this event is concurrent with that of the non ILS related bias in alternative topologies determined by the chi-squared analysis. The non-independence caused by the higher alternative topology (bottom quartet; Figure 1) points to introgression/hybridization between a lineage sister-group to the African (contributing to 31.9% of genes; Figure S4a) or Asian (contributing to 27.9% of genes; Figure S4b) pangolin clades and the ancestor of South-East Asian pangolins (including *M. sp*.). The difference between the two potential basal lineage donors is due to whether we grouped all individuals from a species as the same species (basal Asian contributor), or kept them as separate evolutionary units (basal African contributor). Based on our average divergence estimates (Figure 2; see below), this event likely occurred during the Miocene (between 7.16–16.84 Ma).

**Figure 2:**
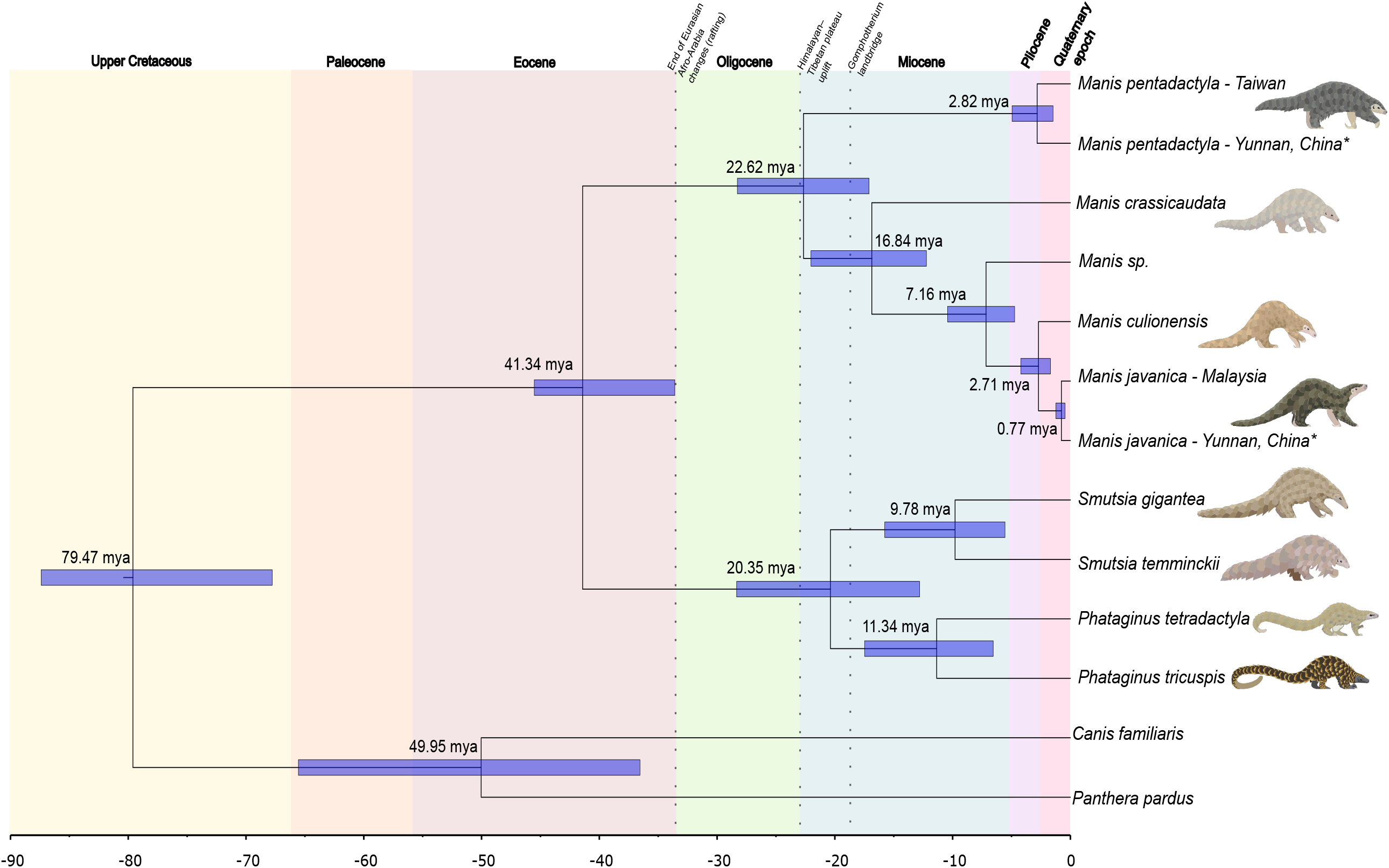
Divergence time estimates along the pangolin phylogenetic tree. Phylogeny with mean posterior divergence times in millions of years and 95% Highest Posterior Density (HPD) intervals (blue bars) for each node (see Table S4 for values). The divergence estimates were based on the unpartitioned, IUPAC concatenated tree dataset of 58 724 014 bp (Figure S3a) using Markov chain Monte Carlo (MCMC) sampling of 20 million generations (2 million generation burn-in) following an auto-correlated, log-normal relaxed clock model. Prior to this, we followed dos Reis and Yang (2019) by first identifying the substitution rate (rgene gamma) for the dataset using baseml and then doing the approximate likelihood calculation by obtaining gradient (g) and Hessian (H) values of branch lengths. We used fossil calibrations on the nodes of Ferae (upper bound from molecular data by Zhou, et al. (2011)), Carnivora and Pholidota as priors for the analysis (Table S3). The various epochs and the Quaternary period are delineated by colored partitions with their corresponding labels above the phylogeny (dates taken from https://stratigraphy.org/timescale/). Three geological/biological events of importance in our study are also highlighted by dotted lines and labels above the phylogeny. Asterisks (*) indicate confiscated individuals whose origins could not be verified. Pangolin illustrations by Sheila McCabe.

### An updated biogeographic scenario for pangolins

Given the uncertainty of the basal branch contributor (Asian or African) in unison with the timing of the admixture event, we can hypothesize that this ghost lineage was likely European. It may have moved towards South-East Asia due to the shrinking of tropical environments in Europe during middle Miocene climatic cooling (Jiménez-Moreno 2006), as seen with hominoids (Begun, et al. 2012). This is plausible as pangolins are suggested to have occurred in Europe within the Miocene (*Necromanis*; Alba, et al. 2018) and into the Pleistocene (S. olteniensis, 2.2–1.9 Ma; Terhune, et al. 2021). Additional population based analyses and fossils across the region will aid in determining whether the hypothesis holds, and whether this ancient admixture event was adaptive (Figueiró, et al. 2017). Recent fossils of pangolins found in South Africa (5 Ma, *S. gigantea*), India (Pleistocene, *M. lydekkeri*) and Java (42 000–47 000 ya, *M. paleojavanica*) provide further evidence of dramatic historical distribution changes (Gaudin, et al. 2009; Terhune, et al. 2021), and at the same time that the potential effect of ghost lineages on our results needs to be considered.

To delve further into the scenario of pangolin diversification, the orthologous whole-genes were used to perform Bayesian estimation of divergence times guided by the coalescent tree and fossil calibrations (Table S3; dos Reis and Yang 2019). At most nodes, our genome-wide results concur with the divergence estimates and resultant biogeographic scenario of diversification previously described with fewer markers (Figure 2; Table S4; Gaubert, et al. 2018; Hu, Roos, et al. 2020). This is the case of the *Manis sp*. taxon which split from *M. culionensis* and *M. javanica* around the late Miocene (7.16 Ma; 4.73–10.42 Ma), similar to the times suggested by Hu, Roos, et al. (2020) using *COI* and *Cytb* markers (6.95 Ma; 4.64–9.85 Ma).

We found that the two South-East Asian pangolin species (*M. javanica* and *M. culionensis*) split during the Upper Miocene to Pliocene (mean = 2.71 Ma; 95% HPD = 1.70–4.21 Ma; Table S4), suggesting that the isolation by sea level rising (800–500 ka) of proto-Philippine pangolins coming from Borneo may have not been the original cause of their divergence (see: Gaubert and Antunes 2015). The split occurs around the same time as the one of the *M. pentadactyla* population in Taiwan and the *M. pentadactyla* individual confiscated in Yunnan, China (2.82 Ma; 1.48–4.95 Ma), and overlaps with the split between the *M. javanica* population in Malaysia and the *M. javanica* individual confiscated in Yunnan, China (0.77 Ma; 0.46–1.24 Ma). These two novel, population-based estimates suggest an alternative hypothesis whereby divergences between lineages through relatively ancient population structuring for *M. javanica, M. culionensis*, and *M. pentadactyla* occurred before isolation of islands, as inferred in the deep, nearly species-level divergence of the two lineages of leopard cats (*Prionailurus bengalensis*) from Sundaland and mainland Southeast Asia (Luo, et al. 2014). This is evidenced by the population divergence of *M. pentadactyla* having occurred even before the final separation of Taiwan from the continental mainland (Kawamura, et al. 2016).

Our analysis indicates more recent divergence time estimates for Asian species than previously suggested (Table S4; Gaubert, et al. 2018), particularly for the genus *Manis* (split between *M. pentadactyla* and the ancestor of *M. crassicaudata*/*M. sp*.*/M. culionensis*/*M. javanica*) which diverged during the Oligocene to Upper Miocene period (22.62 Ma; 17.07–28.23 Ma as opposed to 12.9 Ma; 10.3–15.6 Ma). The split of *M. pentadactyla* likely suggests a northern (*M. pentadactyla*; China) / southern (other *Manis* species; India, Indochina) Asian clade split, coincident with the Oligocene/Miocene boundary’s second uplift of the Himalayan–Tibetan plateau in western China and extrusion of the Indochina block (Spicer, et al. 2020; Deng, et al. 2021). The biogeographic scenarios of other vertebrate species and current distribution limits of *M. javanica* and *M. pentadactyla* across the Ailao Shan-Red River shear zone (Tapponnier, et al. 1990) coincide with this Indochina–China separation (Zhang, et al. 2006; Che, et al. 2010; Xiang, et al. 2021). Overall, more fossil evidence is required to fully comprehend the evolution of pangolins given the paucity of the fossil record along with the possibility of multiple extinct lineages during the evolution of the group (Gebo and Rasmussen 1985; Gaudin, et al. 2009).

### Pangolin demographic history is shaped by lineage-specific biogeography

Inverse instantaneous coalescent rate (IICR) trajectories (Arredondo, et al. 2021), which can be related to fluctuations of effective population size and connectivity, were estimated from the whole genome mapping data of each pangolin species using the pairwise sequentially Markovian coalescent (PSMC) model (Li and Durbin 2011). Although all species showed differing trends, most trajectories started to decline between 1200–600 ka (Figure 3) during the Mid-Pleistocene Transition (Chalk, et al. 2017). This period relates to the transition from the less extreme shorter 41 ka glacial-interglacial cyclicity to that of the more extreme 100 ka (Chalk, et al. 2017), which may have led to population declines or structuring in pangolins as their habitats shifted in size and connectivity (Hewitt 2000; Palkopoulou, et al. 2015). For the Asian species particularly and *P. tetradactyla*, the most rapid phase of the declines occurred during a period that saw one of the two most extreme glaciation periods in the last 800 ka, the Marine Isotope Stage (MIS) 16 (676–621 ka; Lang and Wolff 2011).

**Figure 3:**
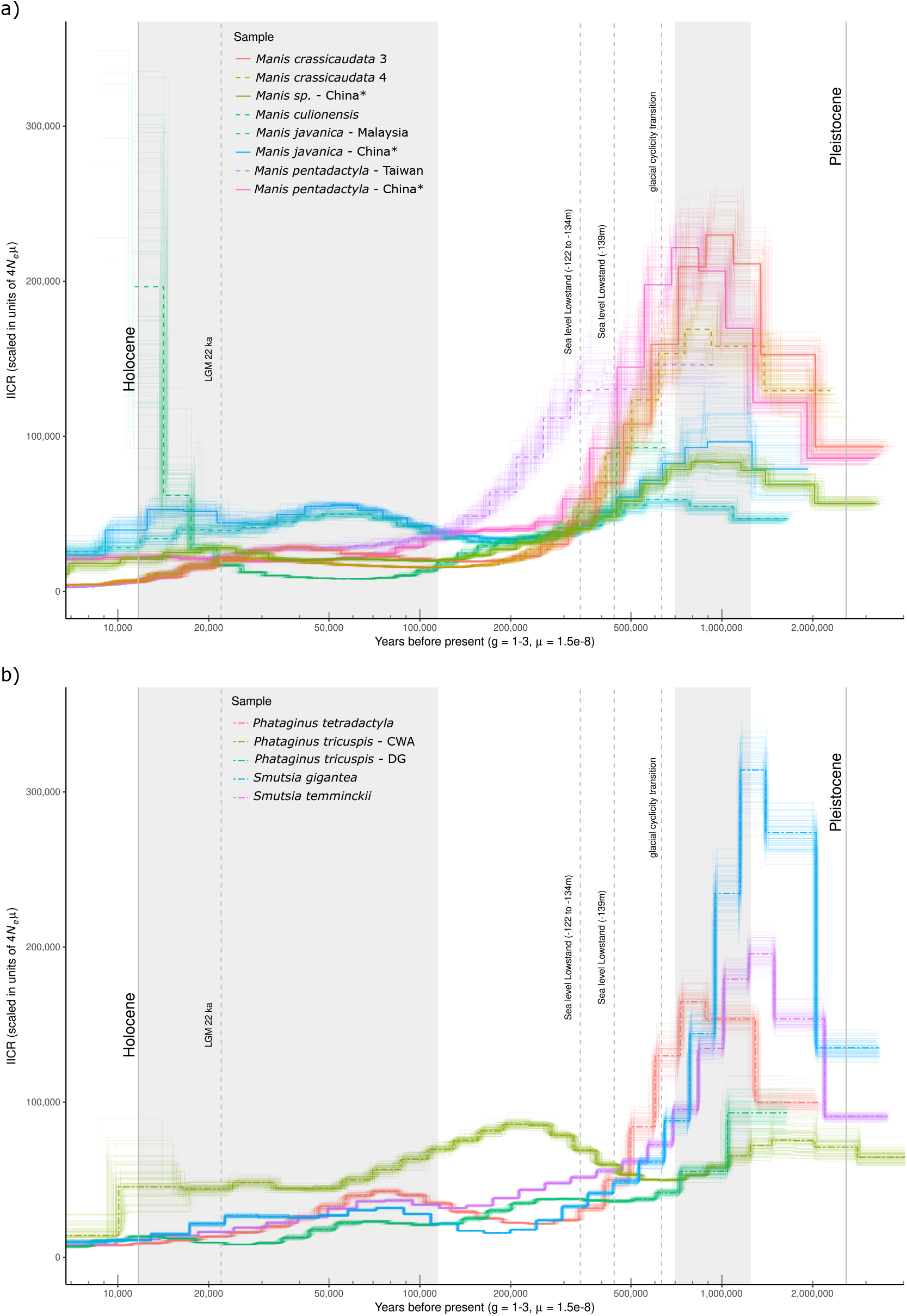
Recent demographic history of pangolin species and populations. Pairwise Sequentially Markovian Coalescent (PSMC) model on diploid genomes provide demographic trajectories of each pangolin species/population for the (a) Asian and (b) African continental clades. Bootstrap support of 100 iterations for each species/population are indicated by the lighter lines of the corresponding species/population colour. Approximate inverse instantaneous coalescence rate (IICR) values are indicative of effective population size (Ne). Curves, however, may be influenced by changes in population size, connectivity and selection. Curves are scaled by a mutation rate of 1.5 × 10^−8^) substitutions per site per generation (µ) based on previous pangolin-related estimates (Choo, et al. 2016). Generation time in years (g) per species was estimated from available literature (Table S6). Vertical dotted lines with labels indicate important climatic events for this study. This figure was produced through a modified version of the following R script: https://github.com/elhumble/SHO_analysis_2020. Asterisks (*) indicate confiscated individuals whose origins could not be verified.

Interestingly, the island species *M. culionensis* (Palawan Isl., Philippines) and the island population of *M. pentadactyla* (Taiwan) exhibited more recent declines (440–300 ka; Figure 3a) post-dating the last two major sea level lowstands of the past 500 ka (Rohling, et al. 1998; Robles, et al. 2015). These lowstand periods [440 ka (−139 m) and 340 ka (−122 – −134 m)] align with fossil and phylogeographic evidence of faunal migration to the current Asian islands during the Middle Pleistocene, including between the strait connecting China to Taiwan (c. 70 m) and the strait connecting Borneo to Palawan (c. 145 m; Tougard 2001; Hosoda, et al. 2011; Kawamura, et al. 2016; Ali 2018). It is therefore possible that after these lowstands, pangolins on these islands were isolated (population structuring) with a decreasing landmass (population decline) as sea levels rose. This difference between island *vs*. continental population fluctuations through time, and the influence of sea-levels for island populations have also been found in South-East island populations of *M. javanica* (Hu, Hao, et al. 2020). The absence of such a signature in the two *M. crassicaudata* individuals from the island population of Sri Lanka could likely be due to the strait connecting India to Sri Lanka being only a minimum of 20 m deep, allowing for extensive continental exchanges across late quaternary climatic fluctuations (Bossuyt, et al. 2004; Ali 2018) as previously suggested from the non-monophyly of Sri Lankan mitogenomes in the species (Gaubert, et al. 2018).

The PSMC curves for the two *P. tricuspis* lineages showed different trends; the Western Central Africa (WCA) lineage presented a stable IICR through time, while the Dahomey Gap (DG) lineage experienced a progressive decline (Figure 3b). These differences are mirrored by their genetic diversity whereby DG contains low levels of genetic diversity and high levels of inbreeding, while WCA has an opposite pattern (Figure 4, see below; Aguillon, et al. 2020; Zanvo, et al. 2022). Our results for the two different *P. tricuspis* lineages (Gaubert, et al. 2016) included in this study suggest that the demographic history of the species is likely complex, lineage specific, and will require more thorough investigation (notably since the DG sample had limited sequencing depth; Table S1). They also confirm that these two lineages have distinct evolutionary trajectories, which have been separated by strong biogeographic barriers (Gaubert, et al. 2016), and thus should be treated as separated conservation units. Further deciphering of phylogeographic structure within pangolin species through population stratification methods is required before more complex demographic scenarios are assessed (Excoffier, et al. 2021). Particularly since the PSMC’s assumption of panmixia and its sensitivity to population structure strongly limit its interpretation (Salmona, et al. 2017; Arredondo, et al. 2021), especially in pangolins, which show high levels of cryptic diversity (Zhang, et al. 2015; Gaubert, et al. 2016; Nash, et al. 2018).

**Figure 4:**
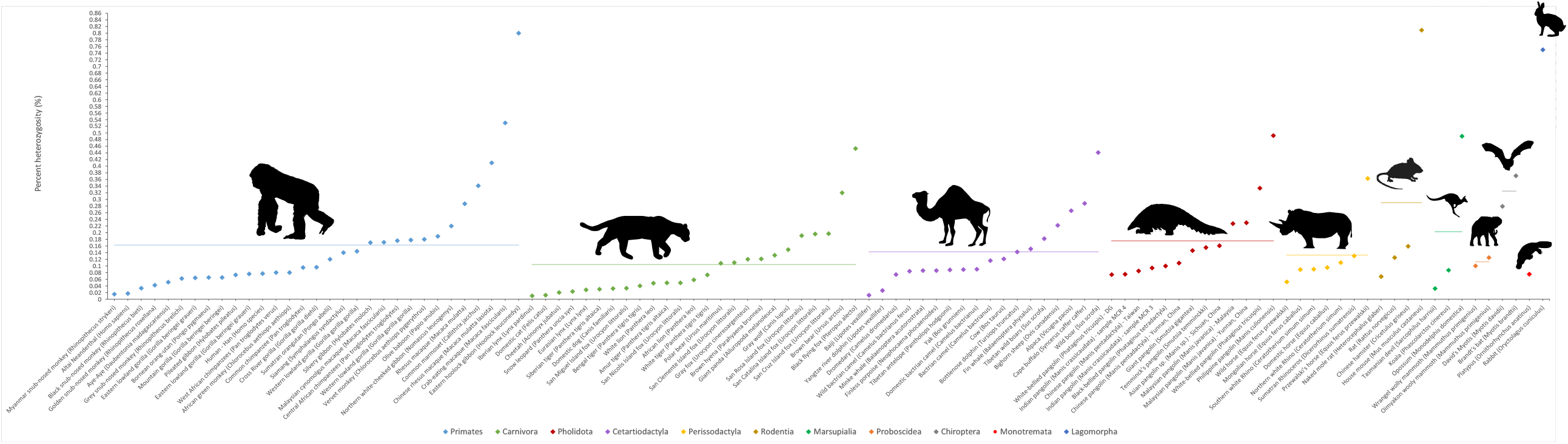
Genome-wide diversity of pangolins within mammalian orders Proportion of genome-wide heterozygosity of pangolins (Pholidota) and other mammals. Estimates are grouped and color-coded by mammalian order with the horizontal line indicating the average proportion of heterozygosity for each order. Estimates are updated from the summary by Hu, Hao, et al. (2020), and displayed from smallest (least diverse) to largest (most diverse) in Table S5. Illustrations indicate the various taxonomic orders and are credited as follows (https://creativecommons.org/licenses/by/3.0/): Pholidota/Certartiodactyla (Steven Traver), Carnivora (Gabriela Palomo-Munoz), Chiroptera (Margot Michaud), Lagomorpha/Marsupialia/Monotremata (Sarah Werning), Perissodactyla (Oscar Sanisidro), Primates (T. Michael Keesey), Proboscidea (Margot Michaud), Rodentia (Jiro Wada) sourced from PhyloPic (http://phylopic.org/).

### Delayed loss in genome-wide diversity

We estimated genome-wide heterozygosity using the site frequency spectrum inferred from genotype-likelihoods of full genome data (Korneliussen, et al. 2014). We found large variation across pangolin species, up to a 6.65 fold divergence between the lowest and highest estimates (0.074 – 0.492% heterozygosity; Figure 4 & Table S5). The *Manis sp*. individual (0.161%) has an above average heterozygosity estimate for mammalian species (average = 0.156%). This taxon is still however comparable to species of conservation importance like the silvery gibbon (*Hylobates moloch*; Table S5). The lower end of the heterozygosity spectrum is occupied by the *P. tricuspis* DG lineage (0.074%), one *M. crassicaudata* individual from Sri Lanka (MCR4; 0.075%), and *M. pentadactyla* (0.085%) from Taiwan. The latter population is from a deeply isolated island, suggesting that the founding effect, isolation from continental populations, and/or subsequent drift have reduced its levels of heterozygosity. This is a concern because long-term low population size and isolation can result in genetic load due to the accumulation and fixation of deleterious variants (Robinson, et al. 2016). However, the major islands hosting pangolin populations (e.g. Sri Lanka, Taiwan, and Borneo) are large when compared to those of iconic examples of such island effect (Robinson, et al. 2016). Therefore, we believe that international trade, habitat fragmentation and finite population structuring through geographical and anthropogenic barriers will probably play a larger role in decreasing heterozygosity levels of these island populations in the future, than their geographic isolation (Lane-deGraaf, et al. 2014).

We hypothesize that the low heterozygosity of the *P. tricuspis* DG lineage can be related to the Dahomey Gap’s bioclimatic history, which shaped a suboptimal savannah habitat intermixed with forest patches. Forest fragmentation through Holocene climatic oscillations (Salzmann and Hoelzmann 2005; Demenou, et al. 2016) in conjunction with high levels of inbreeding through a steady IICR decline over the last 1 million years (Figure 3) may have resulted in a drastic decrease of effective population size and loss of heterozygosity, as noted using microsatellite data (Zanvo, et al. 2022). Along with *Manis culionensis* (0.492%), the other lineage of *P. tricuspis* (WCA; 0.334%) is an outlier for high levels of heterozygosity (Figure 4), alongside common species like the wild boar (*Sus scrofa*) and brown bear (*Ursus arctos*; Table S5). We nevertheless caution that our results may be influenced by genome quality or coverage (as is the case for *M. culionensis*; Mondol, et al. 2013) and that they may not reflect species-wide diversity estimates as they are from only one or two representatives of each species.

Overall, we suggest that conservation management plans should consider the genetic variability and demographic history within pangolin species, with special attention being placed on populations in suboptimal/fragmented habitats (Dahomey Gap) and islands (Taiwan, Palawan, and Sri Lanka). These heterozygosity estimates are likely not yet impacted by the recent boom in illicit pangolin trade (2016 for African species), the biggest contributor to population declines (Mondol, et al. 2013; Challender, et al. 2020). This seems to be the case even for Asian pangolins for which the global trade started a few pangolin generations earlier (Challender, et al. 2020). We therefore suggest that the uninterrupted, global trafficking of pangolins may result in large decreases in heterozygosity estimates with deleterious consequences on the survival ability of pangolins. Such prediction further emphasizes the crucial role that the genomic monitoring of pangolin populations will have to play in the conservation of the Pholidota in the near future.

### Providing genomic resources for pangolin conservation

In light of the drastic loss in genomic diversity likely to come (Dufresnes, et al. 2018; van der Valk, et al. 2019) and no evidence of a reduction in trade (Challender, et al. 2020), the need for genomic data and genetic markers for pangolin conservation is evident. We therefore provide a database of 2 623 genes ranked by levels of mean pairwise identity (Database S1: https://doi.org/10.5281/zenodo.7517409), which could be used to design forensic markers for trade monitoring and prosecution by providing evidence of species traded and possible geographic origin of poaching from seized samples (Zhang, et al. 2015; Gaubert, et al. 2018; Kotze, et al. 2020; Ewart, et al. 2021). Most genes have the ability to discriminate between, and likely within, the eight extant pangolin species. After removing genes that we determined to be outliers (610 genes based on the mean and deviation of mean pairwise identity; Figure S5), the database has 2 623 orthologous genes totaling 3 410 610 segregating/polymorphic sites. The top gene (*TMEM38B*) has a mean pairwise identity of 79.51± 13.56% (min = 65.95%; max = 93.07%) and 20.86% parsimony informative sites. The outlier test is a conservative and basic measure of determining the top genes and so we provide the full list of genes for more in-depth forensic marker discovery. We also provide a good quality reference genome for *S. gigantea* and the draft genomes of pangolin species for which virtually no current genomic data exist. These genomic resources and the resultant inferences we made will be key for better conservation management, further conservation genetic research, and combating the illegal trade of pangolins (Allendorf, et al. 2010; Kotze, et al. 2020; Kardos 2021).

## Acknowledgments

We thank the team at PANGO-GO (https://anr.fr/Project-ANR-17-CE02-0001) for the guidance, provision of material and financial resources to conduct this research. Our gratitude to the following institutions and staff that provided the necessary equipment, resources and guidance: B2M staff at EDB, Get-PlaGe and Genotoul at INRAE - Toulouse, Tshwane University of Technology, the University of Pretoria, and Kalahari Oryx Private Game Farm. We thank Jayanthi Alahakoon for providing samples from Sri Lanka, and Yan Zhuang, Xin Sun, Ke Liu, and Yeyizhou Fu for technical assistance. Our gratitude to Jeremy Johson and the BROAD Institute for providing us with early access to the *Phataginus tricuspis* raw genomic data (GCA_004765945.1). We thank the DNA Zoo (http://www.dnazoo.org) for making their pangolin genomic assemblies publically available. Many thanks to Sheila McCabe for creating and providing illustrations of each pangolin species. This work has been supported by grants from the Agence Nationale de la Recherche (PANGO-GO project: ANR-17-CE02-0001), European Research Council (ConvergeAnt project: ERC-2015-CoG-683257), Mohamed bin Zayed Species Conservation Fund (project: 0925713), the National Research Foundation of South Africa (grant: 71454) and the National Natural Science Foundation of China (NSFC 32070598).

## Author Contributions

P.G. and S.P.H. conceived the study. S.-J.L., F.N., P.G. and D.W.P. collected the samples. P.G., H.M, S.P.H., and M.-K.T., fulfilled laboratory procedures. P.G., S.-J.L., A.D.S.B. and F.D. undertook and funded the genome sequencing. R.A. assembled the *S. gigantea* genome. R.A. and C.S. undertook the annotation and the extraction of CDSs for the three reference genomes. S.P.H. and R.A. undertook draft genome assemblies. S.P.H. performed mapping and variant calling, creation of consensus genes and their alignments, and all the genomic analyses of the study with significant insight from J.S. R.A., M.-K.T. and F.D. wrote the methods section for the *S. gigantea* assembly and annotation. S.P.H. wrote the rest of the manuscript and its supplementary information as well as created the figures and tables, with input from all authors.

## Methods and Materials

### Sample acquisition

Pangolin samples that were used for DNA extraction were collected in the field or museum repositories (see Table S1 for more information). Tissue samples were taken from two deceased *Manis crassicaudata* individuals that were confiscated from the trade in Sri Lanka. Tissue samples from *Phataginus tetradactyla* and *Smutsia gigantea* were collected from the Yaoundé bushmeat market, Cameroon. A spleen sample of *Smutsia temminckii* was collected from an individual that had succumbed to its injuries on an electrified game fence in the Kalahari Oryx Game Farm, South Africa. Finally, we collected a skin sample from *Manis culionensis* at the Field Museum of Natural History, Chicago (FMNH 62919), that had originated from Casuyan, Palawan Island, Philippines. Since the museum sample showed signs of degradation, we took precautions during the mapping and consequent filtering of its short read data as well as interpretation of the results surrounding this species.

### Genome sequencing and assembly

#### *Reference genome of the giant pangolin (*Smutsia gigantea)

High molecular weight DNA suitable for Oxford Nanopore Technologies (ONT) long read sequencing was extracted from an ethanol-preserved muscle sample of a giant pangolin collected in Cameroon (CAM011) using the protocol optimized by Tilak, et al. (2020). Long read sequencing was performed with the ONT MinION instrument using libraries prepared with the ONT Ligation Sequencing kit SQK-LSK109 on four flow cells (FLO-MIN-106). This generated a total of 31.7 Gb of raw long read data representing a genome depth of coverage of about 13x. ONT raw signal FAST5 files were then base-called with the high accuracy mode of Guppy v3.2.4 (https://github.com/nanoporetech) on a GPU computing server of the Montpellier Bioinformatics Biodiversity platform (https://mbb.univ-montp2.fr/MBB) and subsequently cleaned using Porechop v0.2.3 (https://github.com/rrwick/Porechop). Complementary Illumina 150PE short reads were produced on an Illumina HiSeq 3000 sequencing system at the GeT-PlaGe sequencing platform at Genotoul (https://www.genotoul.fr). After cleaning using Trimmomatic v0.33 (options -phred33 LEADING:3 TRAILING:3 SLIDINGWINDOW:4:15 MINLEN:50; (Bolger, et al. 2014)), we obtained about 185 Gb of short read data representing a genome depth-of-coverage of about 87x (Table S1). A *de novo* hybrid genome assembly was performed using MaSuRCA v3.2.9 (Zimin, et al. 2013), which first assembles short reads into “super-reads” before assembling them guided by the long ONT reads. To evaluate genome quality, traditional measures like the BUSCO score (Waterhouse, et al. 2018), the number of scaffolds and contig N50, and mean and maximum lengths were computed.

*De novo* genome annotation was performed for our *S. gigantea* hybrid assembly and the DNA Zoo chromosome-length assembly of *P. tricuspis* following the approach used in Allio, et al. (2021). First, repetitive elements were annotated and masked to avoid producing false evidence for gene annotations (Yandell and Ence 2012). This annotation was first performed for each genome independently using RepeatModeler v2.0.2 (Smit and Hubley 2008). To improve the accuracy of these *de novo* annotations, the libraries obtained from the different genomes were cleaned by removing protein-like sequences and were clustered for further analyses. The second step was to identify repeat elements by similarity search against publicly available libraries of mammalian repeats (DFAM (Wheeler, et al. 2012)) using RepeatMasker v4.1.2-p1 (Tarailo-Graovac and Chen 2009). The annotations resulting from these two steps were synthesized in a GFF file to be fed into MAKER v3 (Holt and Yandell 2011).

To improve the gene annotation, we relied on transcriptomic information using publicly available RNAseq data for *M. javanica* (Table S1). To do so, raw reads from 25 transcriptomes were downloaded from SRA, cleaned using fastp v0.20.0 (Chen, et al. 2018) using default parameters, and assembled with Trinity v2.9.0 (Grabherr, et al. 2011). The resulting transcriptome assemblies were annotated with an adapted version of assembly2ORF (https://github.com/ellefeg/assembly2orf), which was specifically designed to annotate transcriptomes. This pipeline relies on evidence-based gene predictions to extract and annotate gene CDSs from transcriptome assemblies. The CDSs resulting from the annotation were concatenated and clustered by similarity with CD-HIT v4.8.1 (Li and Godzik 2006) to improve the efficiency of the subsequent MAKER annotation. The CDSs obtained from RNA-seq data were fed into MAKER v3 to help the evidence-based gene prediction. Additionally, the manually annotated, non-redundant protein sequence database Uniprot/SWISSPROT (Bairoch and Apweiler 2000; The UniProt Consortium 2009) was provided to MAKER v3 for the annotation.

To improve the annotation, three runs of MAKER v3 were performed iteratively. In the first run, evidence-based gene predictions using sequence similarities with the CDSs extracted from both the Swiss-prot database and the transcriptomes were computed. Then, two additional runs were performed, with SNAP v2006-07-28 (Korf 2004) and Augustus v3.2.3 ((Stanke, et al. 2006); via BUSCO v3, (Waterhouse, et al. 2018)) as implemented within MAKER v3 to help create more sound gene models. In doing this, MAKER v3 uses the annotations from the two prediction programs in addition to the evidence-based gene predictions (similarities with reference CDSs) when constructing its models.

#### Draft genomes for other pangolin species

Paired-end Illumina short read (150 bp) sequencing was conducted on *Manis crassicaudata, M. culionensis, S. temminckii* and *P. tetradactyla* samples ranging from 44x to 76x sequencing depth (Table S1). SOAPdenovo2 vr240 (Luo, et al. 2012) was used to assemble the genomes of all sequences generated in this study. The best kmer length assembly for each species was identified using SeqKit v0.9.3 (Shen, et al. 2016) statistics on the scaffolds (N50, mean and maximum length and the assemblies were gap-closed with GapCloser v1.12 (SOAPdenovo2).

#### Additional genomic data

Short read data from two previously published genomes of *M. javanica* and *M. pentadactyla* (Choo, et al. 2016; Hu, Hao, et al. 2020), an uncertain *Manis* species (labeled as *M. crassicaudata* on NCBI but published as *M. culionensis*) (Cao, et al. 2021), and one current draft genome of *P. tricuspis* were extracted from the Sequence Read Archive (SRA) on NCBI using the SRA toolkit v2.9.6 (Table S1; Leinonen, et al. 2010). We also obtained chromosome-scale genome assemblies of *P. tricuspis* (https://www.dnazoo.org/assemblies/Phataginus_tricuspis; Choo, et al. 2016) and *M. javanica* (https://www.dnazoo.org/assemblies/Phataginus_tricuspis) from DNA Zoo (Dudchenko, et al. 2017; Dudchenko, et al. 2018). The former also included short read data that we used in our analyses (Table S1).

### Single-copy orthologous gene dataset

To build a comprehensive phylogenomic dataset for all extant pangolin species, we relied on the OrthoMaM v10 database (Scornavacca, et al. 2019), which is composed of 14 509 single-copy orthologous genes for 116 mammal species, including 13 403 CDSs (coding DNA sequences) for *M. javanica*. We implemented a pipeline in which we used a dual strategy to reduce the effect on the high level of divergence between pangolin species (particularly between Asian and African clades) if one reference were used. First we used the *de novo* annotations to conduct single-copy orthologous gene extraction of the DNA Zoo reference genome assemblies for each pangolin clade (Manis javanica for Asian clade and Phataginus tricuspis for African clade). This provided us with what we termed reference CDSs and their IDs. This was followed by subsequent mapping of Illumina reads of all species on their clade-specific reference genome assemblies and using the IDs from the CDS references to extract the relevant orthologous genes (Figure S1).

#### *CDS extraction from* de novo *reference assemblies*

To extract the CDSs specifically corresponding to the single-copy orthologs of the OrthoMaM database, for each orthologous gene alignment, a HMM profile was created via *hmmbuild* of the HMMER toolkit v3.1b2 (Eddy 2011). Then, all HMM profiles were concatenated and summarized using *hmmpress* to construct a HMM database. Finally, for each CDS newly annotated by MAKER v3, *hmmscan* was used on the HMM database to retrieve the best hits among the orthologous gene alignments. For each orthologous gene alignment, the most similar sequences for each species were detected via *hmmsearch*. Outputs from *hmmsearch* and *hmmscan* were discarded if the first hit score was not substantially better than the second (hit 2< 0.9 hit 1). This ensured that our orthology predictions for the newly annotated CDSs were robust.

From this and previous work, we had obtained orthologous CDS information from *Manis* (*M. javanica* from GenBank annotation and CDS extraction from OMM), *Phataginus* (*P. tricuspis* from DNAZoo, annotation and CDS extraction from this study), and *Smutsia* (*S. gigantea*, annotation and CDS extraction from this study).

#### Genome-wide mapping and consensus

Short read data from all species were cleaned with fastp v0.19.4 with a base pair quality threshold (>20 Phred) and the paired-end base correction function using overlapped reads (-c option). The cleaned reads were then mapped to their respective clade-specific reference genome assemblies (Asian species: *M. javanica* from GenBank; *Phataginus*: *P. tricuspis* from DNA Zoo) using the default settings of the BWA-MEM algorithm of BWA v0.7.15 (Li and Durbin 2009) after testing mismatch, clipping and gap penalties. However, due to the lower quality of the sequencing data of *Manis culionensis*, we applied a more stringent mapping approach (seed = 23, mismatch penalty = 7). We used SAMtools v1.10 (Li, et al. 2009) *view* keep only reads mapped into proper pair (-f 2) and mapped reads with a high mapping quality (>30 Phred). These filters were tested using SAMtools v1.10 flagstat before and after filtering, and using Qualimap 2 v0.7.1 (Okonechnikov, et al. 2015) after filtering. Resulting bam files were sorted with SAMtools v1.10 *sort*, duplicates marked with Picard v2.20.7 *MarkDuplicates*, as well as genome-wide mean depth and proportion covered at a depth of >1x and 10x that were calculated using SAMtools v1.10 *depth* and a custom script (Custom script 1). ANGSD v0.933 (Korneliussen, et al. 2014) *dofasta* - was used to obtain consensus fasta sequences files for each bam file using genotype likelihoods. Both option 3 (most common allele chosen to make haploid consensus) and option 4 (multiple alleles chosen to make IUPAC consensus) were chosen in conjunction with command *docounts* 1. Reads were filtered if they were of bad quality (-remove_bads), not paired (-only_proper_pairs), below a mapping quality of Phred 30 (-minmapq), and had multiple mappings (-uniqueOnly). We also filtered for bases below Phred 20 (-minQ; except *M. culionensis* which had Phred 30) and a minimum sequence depth of 10x (-setMinDepth) and a maximum of 2 times that of the average depth per individual (-setMaxDepth). Outgroup taxa *Panthera pardus* (GCF_001857705.1) (Kim, et al. 2016) and *Canis familiaris* (https://www.dnazoo.org/assemblies/Canis_lupus_familiaris_Basenji), which are part of the sister order Carnivora (Liu, et al. 2017), underwent the same process with each being mapped to their respective Hi-C reference genomes from DNAZoo (Dudchenko, et al. 2017; Dudchenko, et al. 2018).

#### Obtaining orthologous full gene markers using orthologous CDS IDs

Using the orthologous CDS gene IDs from OrthoMaM for each pangolin reference (*M. javanica* and *P. tricuspis*), whole genes annotations containing these IDs were extracted from the annotation files of the references (*M. javanica, P. tricuspis* and the two outgroup species; Custom script 2). Annotations with duplicates and those that were not found in both pangolin references were removed. This left us with gff3 annotation files of 5 660 orthologous full genes. BEDtools v2.29.0 (Quinlan and Hall 2010) *getfasta* was used to extract these orthologous full genes from the full genome IUPAC and haploid consensuses, after being indexed with SAMtools v1.10 *faidx*.

### Phylogenomics and forensic markers from genes

Multi-gene fasta files of each species were converted into multi-species fasta files per gene marker using SeqKit v0.9.3 (Shen, et al. 2016) *split* (--by-id option) and subsequent concatenation using the *cat* command. These consisted of all eight species (including two representatives each of *M. javanica* and *M. pentadactyla*) and the outgroup taxa from the Order Carnivora. The dataset was then split into two parts, one with haploid consensus genes of only one representative of the eight known species, and the second with IUPAC consensus genes of all aforementioned pangolin individuals and the outgroup taxa. The first was used to determine per-gene diversity estimates across pangolins and the second was used for phylogenetic testing, divergence time estimation, and identifying reticulation events.

#### Genetic diversity estimates from whole genes

Obtaining genetic diversity estimates for pangolins from genes requires the use of one representative per species and the removal of outgroup taxa as these would bias the level of diversity per gene. Hence, the two Carnivora outgroup taxa, together with *M. javanica, M. pentadactyla*, and *M. sp*. (from Choo, et al. (2016); Cao, et al. (2021)) were removed from the gene dataset before undergoing alignment. Sequences were aligned using MAFFT v7.313 (Katoh and Standley 2013) --auto option and [Transitive Consistency Score (TCS)](https://tcoffee.org/Projects/tcs/index.html) (Chang, et al. 2014) for each alignment were assessed using T-COFFEE v11.00.8 (Notredame, et al. 2000) *evaluate* with the clustalw2_msa method. Multiple Sequence Alignments (MSAs) with TCS scores under 80 were removed based on the possibility of paralogy, repetitive regions or mis-annotated genes between the two pangolin reference genomes. This resulted in 3 238 gene MSAs of high quality for further analyses. PhyKIT v1.1.3 (Steenwyk, et al. 2021) was used to obtain statistics on the pairwise proportional identity of each marker (command *pairwise_identity*) and parsimony informative sites (command *parsimony_informative_sites*). Due to PhyKit calculating alignment gaps as informative sites, all gaps were removed prior to this analysis using TrimAL v1.4.1 (Capella-Gutiérrez, et al. 2009) with the option --nogaps. DnaSP v6 (Rozas, et al. 2017) was used to conduct the “DNA Polymorphism” analysis in batch mode to obtain the rest of the diversity estimates. These outputs were then merged with a custom script (Custom script 3) to obtain a range of diversity statistics per gene. Using the package “robustbase” v0.93 (Maechler, et al. 2021) in R v3.6.1 (Rstudio Inc., Massachusetts, U.S.A), we identified possible outlying genes with low levels of mean proportional pairwise identity (or high levels of diversity between species) by filtering values over two Qn deviations from the median of both the mean (487 outliers) and standard deviations (566 outliers) of pairwise identities (Figure S5) (Rousseeuw and Croux 1993). This was done to prevent inconsistent markers being used in forensic screening approaches due in part to possible biological reasons (paralogy, repetitive elements, inconsistency across taxa, etc.).

#### Phylogenomic tree building and concordance analyses

From the results on TCS alignments scores above, the same 3 238 cleaned, orthologous genes with all pangolin individuals and outgroup taxa were aligned and trimmed following the aforementioned protocol (MAFFT v7.313 and TrimAL v1.4.1), however the –gappyout option was used for trimming. Additionally, these alignments stem from IUPAC consensus and they were again evaluated with T-COFFEE v11.00.8 in order to remove outgroup taxa that had a TCS score of lower than 90 (using SeqKit v0.9.3 *grep*). Alignment statistics of each MSA file per gene were obtained and were then concatenated into a single MSA using AMAS v0.98 *summary* and *concat* (Borowiec 2016) with --part-format set to RAxML in order to provide a partitioning file by gene. ModelTest-NG v0.1.5 (Darriba, et al. 2019) was used to identify the best model of sequence evolution for each gene-partition (input partition file from AMAS) as well as the entire alignment (no partition). We set the -h option to uigf (tests rate heterogeneity: Uniform, +I, +G, and +I & +G) and -T option (model test template) to phyml (all 11 models of evolution) in order to test 88 DNA models. Partitioned and non-partitioned concatenated IUPAC phylogenies (supermatrix) with 1 000 Felsenstein bootstrap replicates were inferred from the concatenated MSAs using RAxML-NG v0.9.0 (Kozlov, et al. 2019) following the best-fitted DNA model for each partition under the BIC test. A multiple species coalescent summary tree was also inferred from the MSAs by first using RAxML-NG v0.9.0 to build a gene tree per gene whilst accounting for the best-fitting DNA model (BIC) for each gene (extracted from the aforementioned ModelTest-NG analysis), followed by summarizing these gene trees with ASTRAL-III v5.6.3 (Zhang, et al. 2018). A polytomy test (-t10) using these data was conducted in order to test whether each branch is a potential polytomy (Sayyari and Mirarab 2018), but none were suggested as a polytomy. Finally, gene (--gcf) and site (--scf with 100 quartets randomly sampled around each internal branch) concordance factor analyses based on the aforementioned gene trees and ASTRAL-III output (coalescent species tree) were performed with IQ-TREE 2 v2.0.6 (Minh, Schmidt, et al. 2020). This provides a full analysis of the raw data in terms of genes or sites that may be disagreeing with the species tree phylogeny. This was followed by a Chi-squared test of independence in R v3.6.1 (Rstudio Inc., Massachusetts, U.S.A) based on a script designed by Robert Lanfear (http://www.robertlanfear.com/blog/files/concordance_factors.html). This tests the frequencies of two alternative quartet topologies of each internal branch whereby significance indicates the possibility of something other than incomplete lineage sorting causing gene tree discordance. We did not use this test on site concordance frequencies, as this test does not take linkage disequilibrium between sites into account, which can be assumed to play a large role for multiple sites in a single gene.

#### Reticulate evolutionary relationships

We used PhyloNet v3.8.2 (Than, et al. 2008) to run two separate analyses in order to determine possible reticulation events through maximum pseudolikelihood estimates. The first was to consider each individual as a separate species and the second was to merge multiple individuals of the same species as one. Both analyses were conducted by inputting the RAxML gene trees whilst the best number of reticulation events (0-4) were determined by comparing the change in maximum pseudolikelihood scores between them. PhyloPlot in Julia (Solís-Lemus, et al. 2017) was used to visualize the networks.

#### Divergence time estimates

We used MCMCTree in the PAML-4.9h package (Yang 2007) to estimate divergence times on the non-partitioned RAxML-NG (supermatrix) phylogeny inferred from the IUPAC consensus gene dataset. We used Ferae (root), Pholidota and Carnivora fossils as upper and lower bound calibrations (Table S3). We followed dos Reis and Yang (2019) by first identifying the substitution rate (rgene gamma) for the IUPAC gene dataset using baseml and then doing the approximate likelihood calculation by obtaining gradient (g) and Hessian (H) values for branch lengths. The MCMC sampling from the posterior distribution of times and rates using MCMCTree was run for 20 million generations with a burn-in of 2 million generations and was run twice to test for convergence in Tracer from BEAST 2 (Bouckaert, et al. 2014). An auto-correlated, log-normal relaxed clock model was implemented. Finally, we ran the analysis by sampling from the prior of times and rates (usedata = 0) to test the soundness of the prior and whether our calibrations may influence the results.

### Mitochondrial genes and genomes

To obtain mitochondrial genomes of each species in this study as well as those from the undetermined *Manis sp*. (Cao, et al. 2021) and the two unpublished *P. tricuspis* reference genomes, 10 million reads were extracted from fastq files and mapped to a published mitochondrial genome reference of each species using the default options of Geneious mapper in Geneious v9.1.8 (Kearse, et al. 2012). These were then consensus called and aligned to the mitochondrial genome dataset from Gaubert, et al. (2018) with MUSCLE (10 iterations) in Geneious. The alignment underwent Neighbour-Joining (NJ) phylogeny testing in MEGA X (Kumar, et al. 2018) with Kimura 2-parameter model of evolution and 1000 bootstrap replicates. We also aligned the *COI* and *Cytb* genes from two samples of the potential new Asian species suggested by Hu, Roos, et al. (2020) to the mitochondrial genome dataset and extracted these regions for an additional NJ phylogeny of these two genes.

### Genome-wide analyses

The aforementioned cleaned short read data for each pangolin species were mapped to their closest reference per genus (*Phataginus*: *P. tricuspis* DNA Zoo; *Manis*: *M. javanica* DNA Zoo; *Smutsia*: *S. gigantea* this study) using BWA-MEM from BWA v0.7.15. Resulting bam files were filtered using SAMtools v1.10 *view* (-f 2; >30 Phred), sorted with SAMtools v1.10 *sort*, had duplicates marked with Picard v2.20.7 *MarkDuplicates*, as well as had genome-wide mean depth and proportion covered at a depth of >1x and 10x that were calculated using SAMtools v1.10 *depth* and a custom script (Custom script 1).

#### Comparative genome-wide heterozygosity

ANGSD v0.933 (Korneliussen, et al. 2014) was used to estimate genome-wide heterozygosity per sample using a script adapted from de Jager, et al. (2021) following the ANGSD workflow (http://www.popgen.dk/angsd/index.php/Heterozygosity). To do this, the command *doSaf* was implemented on the alignment bam file of each individual from genome-wide mapping to infer its folded site allele frequency likelihood. We used the genus-specific reference as the ancestral state and the SAMtools method for genotype likelihoods (-GL 1) as it considers possible sequencing errors and performs necessary corrections. The option -C 50 was used to adjust mapping quality for excessive mismatches, and reads were removed if they were of bad quality (-remove_bads), not paired (-only_proper_pairs), below a mapping quality of Phred 30 (-minmapq), and had multiple mappings (-uniqueOnly). Bases below Phred 20 were also removed (-minQ), with the exception of *M. culionensis* which had Phred 30. After *doSaf*, we inferred the site frequency spectrum with the ANGSD sub-program realSFS, and finally, the genome-wide heterozygosity by dividing the number of heterozygous sites by the total number of sites per genome. We extended the list of genome-wide heterozygosity estimates for mammalian species built by (Hu, Hao, et al. 2020), many of which are of conservation importance, by including our estimates as well as additional estimates published since the conception of the table.

#### Demographic history reconstruction with PSMC

Each alignment file (bam) from the whole-genome pipeline output was converted to a diploid consensus fastq file by first determining genotype likelihoods using BCFtools v1.8 *mpileup* (minimum base and mapping quality of Phred 30), calling the genotypes with BCFtools v1.8 *call* (-c option for consensus caller), and then filtering and converting the vcf to a fastq file using the vcftultils.pl vcf2fq script (SAMtools v1.10). Filtering with vcfutils.pl included a minimum mapping quality of Phred 30 (-Q), a minimum coverage threshold of 10x (-d) and maximum coverage threshold of double the mean genome-wide coverage of each individual (-D) as determined from the whole genome pipeline. For all genomes, the mean genome-wide depth was higher than the recommended PSMC cutoff of ≥18x and the percentage of missing data per genome at ≥10x depth was lower than the recommended PSMC cutoff of 25% (Nadachowska-Brzyska, et al. 2016). This was the case except for the one of two *M. crassicaudata* (MCras 4; depth = 9X and 37% of reference covered at ≥10x) and *P. tricuspis* (From DNA Zoo; depth = 10.8X and 45% of reference covered at ≥10x) genomes. These were only used for PSMC and heterozygosity estimates and should be taken with caution.

Using the program PSMC v0.6.5 (Li and Durbin 2011), the fastq consensus files with diploid variant information were then converted to fasta-like files (psmcfa) containing information of whether there is at least one homozygote in a bin of 100 bp. From there, we ran the PSMC analysis following the default settings described by Li and Durbin (2011) for human populations (-N 25, -t 15, -r 5, -p “4+25*2+4+6”) and used on other mammalian species (Westbury, et al. 2018; Chen, et al. 2019; de Jager, et al. 2021). By randomly sampling subsections of the psmcfa file, 100 bootstrap replicate analyses were performed in order to estimate the variance in the approximate inverse instantaneous coalescence rate (IICR). This is the inverse of the rate at which coalescence events take place through time, which was originally used as a function of effective population size in a panmitic model (Li and Durbin 2011). However, inferences must take into account the potential of a non-panmitic model of demographic history and the confounding effects of natural selection and population structure, which can also affect coalescence (Arredondo, et al. 2021; Johri, et al. 2021). A mutation rate of 1.5×10^−8^) years/site was implemented (Choo, et al. 2016), while a generation time in years was chosen for each species depending on available literature (Table S6). The PSMC figures combining the species from each continental clade were constructed in R v5.1 (Rstudio Inc., Massachusetts, U.S.A) following de Jager, et al. (2021) by editing a script (found at: https://github.com/elhumble/SHO_analysis_2020) that utilizes the plotPsmcR function (Liu and Hansen 2017).

## Data accessibility

Draft genomes (*Manis culionensis, M. crassicaudata, Phataginus tetradactyla, Smutsia temminckii*) and a reference genome with associated metadata (*S. gigantea*) have been deposited in Genbank and are publicly available as of the date of publication. The associated sequence read data have also been deposited in Genbank (SRA) for the aforementioned genomes (except for *S. temminckii*). This paper also analyzes existing, publicly available genomic data (*M. pentadactyla, M. javanica, P. tricuspis*). The accession numbers or links for the aforementioned genomic data are listed in Table S1.

A database containing the list orthologous, whole-genes ranked by diversity amongst all eight pangolin species has been deposited at Zenodo and is publicly available (Database S1: https://doi.org/10.5281/zenodo.7517409). All original code in the form of custom scripts for processing the genomics data in this study have been deposited at Zenodo and are publicly available (https://doi.org/10.5281/zenodo.7517409).

## Supplemental information

Document S1. Figures S1–S5 and Tables S1–S6

Database S1. Genes ranked by diversity. Available at Zenodo (https://doi.org/10.5281/zenodo.7517409)

Custom scripts 1-3. Available at Zenodo (https://doi.org/10.5281/zenodo.7517409

