## Supplementary Figures and Tables for "Pangolin genomes offer key insights and resources for the world’s most trafficked wild mammals"

Whole-Gene Pipeline

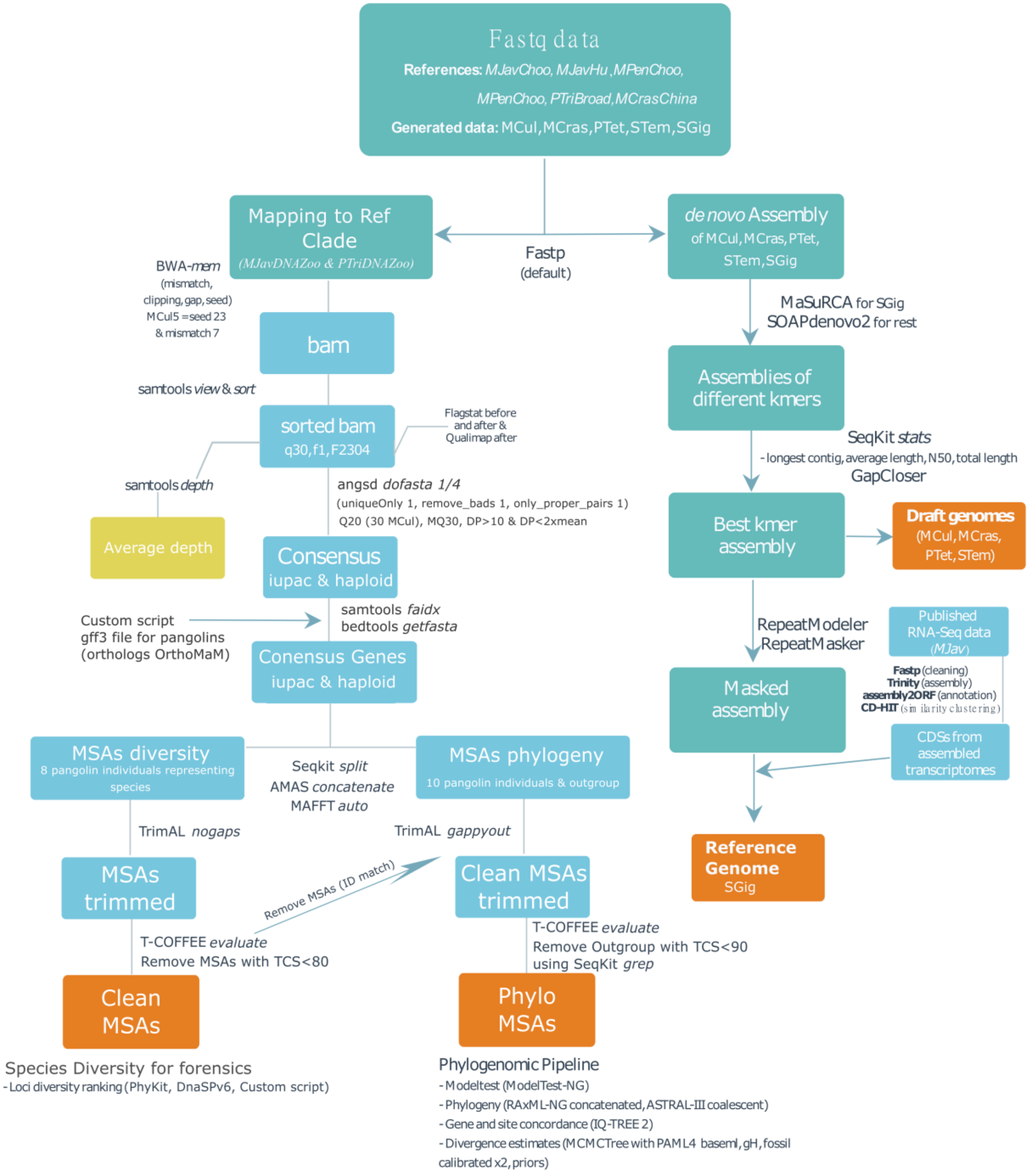


Figure S1: Orthologous whole-gene marker pipeline and subsequent analyses.

This pipeline was formulated and used to create the whole-gene markers (introns and exons) required for further analyses. During this process, the reference genome (*Smutsia gigantea*) and draft genomes for four other pangolin species were created. First, the *de novo* assembly side of the pipeline (right side) was conducted before the whole genome and subsequent whole-gene extraction side of the pipeline could be conducted (left side). Custom scripts can be found at Zenodo (https://doi.org/10.5281/zenodo.7517409). We used shorthand notations to indicate the genomes used for each species as follows: MJavChoo - *Manis javanica* from Malaysia (Choo, et al. 2016), MJavHu – *M. javanica* confiscated in China (Hu, Hao, et al. 2020), MPenChoo – *Manis pentadactyla* from Taiwan (Choo, et al. 2016), MPenHu – *M. pentadactyla* confiscated in China (Hu, Hao, et al. 2020), MCrasChina – *Manis sp.* confiscated in China (Cao, et al. 2021), PTriBROAD – *Phataginus tricuspis* (unpublished BROAD institute; GCA_004765945.1), PTriDNAZoo – *P. tricuspis* (unpublished DNA Zoo; https://www.dnazoo.org/assemblies/Phataginus_tricuspis), MCul – *M. culionensis* (this study), MCras – *M. crassicaudata* (this study), PTet – *P. tetradactyla* (this study), SGig – *Smutsia gigantea* (this study), STem – *S. temminckii* (this study). Further information on these genomes and samples can be found in Table S1.

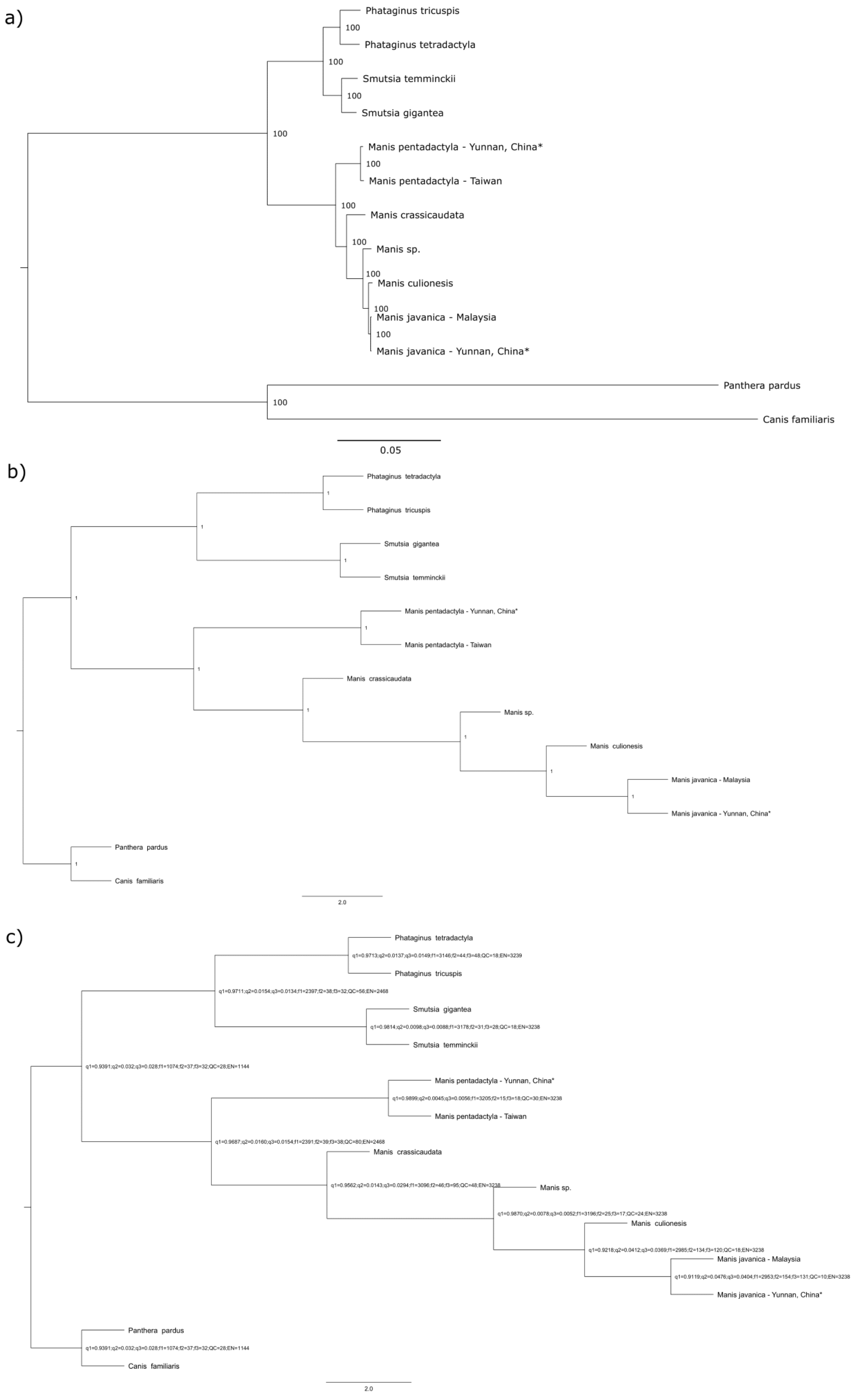


Figure S2: Phylogenomic relationships of pangolins inferred from concatenated and coalescent trees

a) Non-partitioned concatenated phylogeny based on 58 724 014 bp from 2 238 IUPAC consensus whole-gene markers. The pangolin phylogeny consists of 13 individuals from all eight species and is rooted with two representatives of the sister Order Carnivora (*Canis familiaris* and *Panthera pardus*). Nodal values indicate bootstrap support from 1 000 Felsenstein replicates. The model used for the maximum likelihood tree search was the TVM+FO+G4m based on a best-fit DNA model test.

b) Multiple species coalescent phylogeny based on 2 238 gene trees. Branch lengths are in coalescent units while nodal values indicate local posterior probability support (1=complete support).

c) Multiple species coalescent phylogeny based on 2 238 gene trees. Branch lengths are in coalescent units. Nodal values indicate the level of gene tree conflict, which is calculated through the number of alternative gene tree quartets that agree with the main species tree quartet topology (this figure) at each internal branch. The normalized quartet score for the entire phylogeny is 0.981. Using a polytomy test we did not identify any polytomies in the tree (which could influence these results). q1, q2, and q3 refer to the proportion of quartets in the gene trees that agree with a branch (quartet support) for the main topology (LR|SO), first alternative (RS|LO) and second alternative (RO|LS), respectively. f1, f2, and f3 refer to the same as above but are the raw number of quartet trees instead of the proportion. QC is the total number of quartets possible around each branch and EN is the number of effective genes for each branch.

Asterisks (*) indicate confiscated individuals whose origins could not be verified.

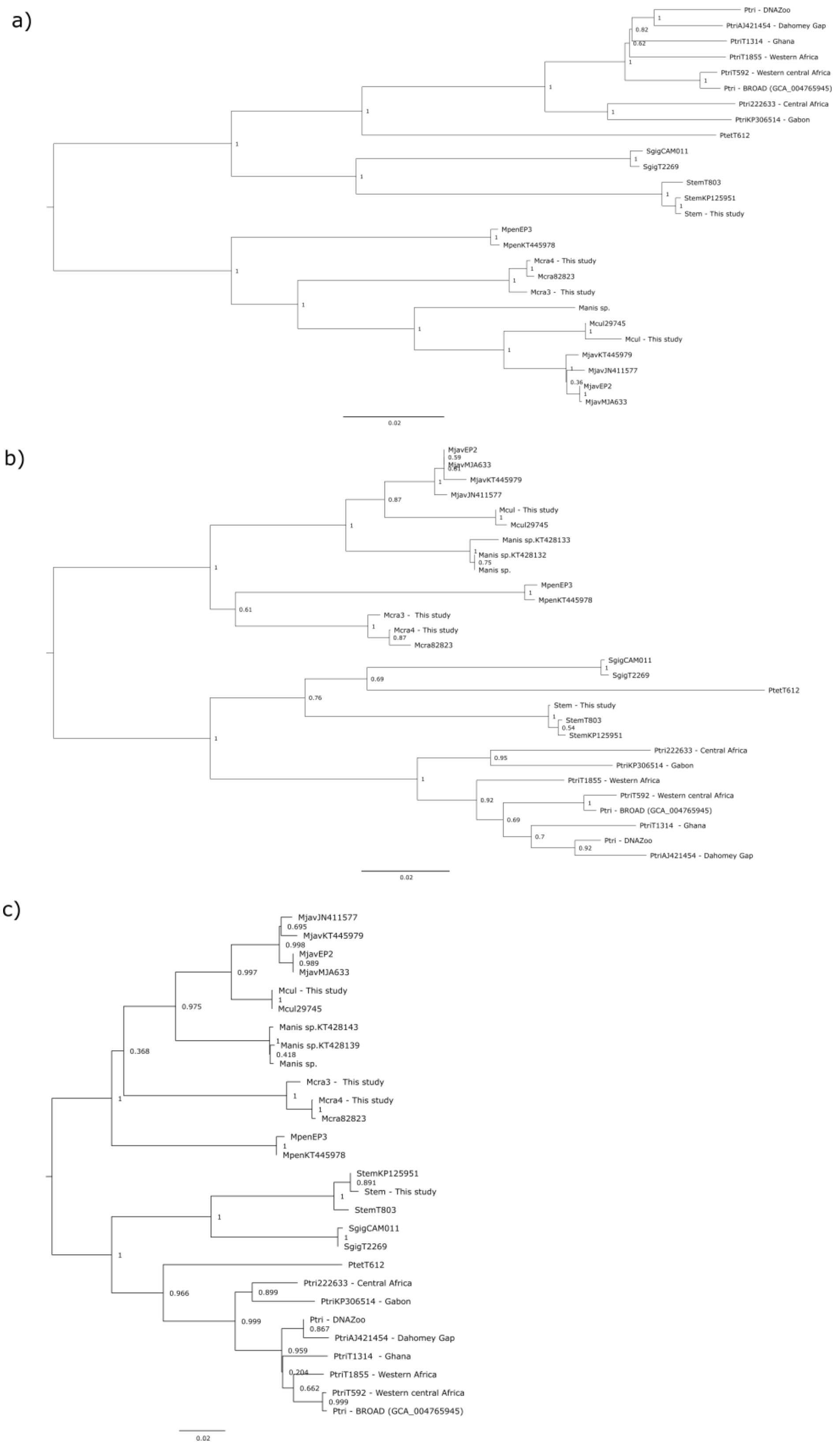


Figure S3: Phylogenetic relationships of pangolins inferred from mitochondrial gene and genome trees

a) Full mitochondrial genome phylogeny based on 16 437 bp using the Neighbor-Joining method in MEGA X (Kumar, et al. 2018). The pangolin phylogeny consists of 26 individuals from all eight species and six cryptic lineages of *Phataginus tricuspis*. Nodal values relate to bootstrap support (1000 replicates). The DNA model of evolution was Kimura 2-parameter method with all positions with less than 90% site coverage eliminated (partial deletion option).

b) Full cytochrome b (*Cytb*) gene phylogeny based on 399 bp using the Neighbor-Joining method in MEGA X (Kumar, et al. 2018). The pangolin phylogeny consists of 28 individuals, including that from all eight species, the six cryptic lineages of *Phataginus tricuspis,* and the two samples suggested as a possibly new *Manis* species (Hu, Roos, et al. 2020). Nodal values relate to bootstrap support (1000 replicates). The DNA model of evolution was Kimura 2-parameter method with all positions with less than 90% site coverage eliminated (partial deletion option).

c) Full cytochrome oxidase subunit I (*COI*) gene phylogeny based on 600 bp using the Neighbor-Joining method in MEGA X (Kumar, et al. 2018). The pangolin phylogeny consists of 28 individuals, including that from all eight species, the six cryptic lineages of *Phataginus tricuspis,* and the two samples suggested as a possibly new *Manis* species (Hu, Roos, et al. 2020). Nodal values relate to bootstrap support (1000 replicates). The DNA model of evolution was Kimura 2-parameter method with all positions with less than 90% site coverage eliminated (partial deletion option).

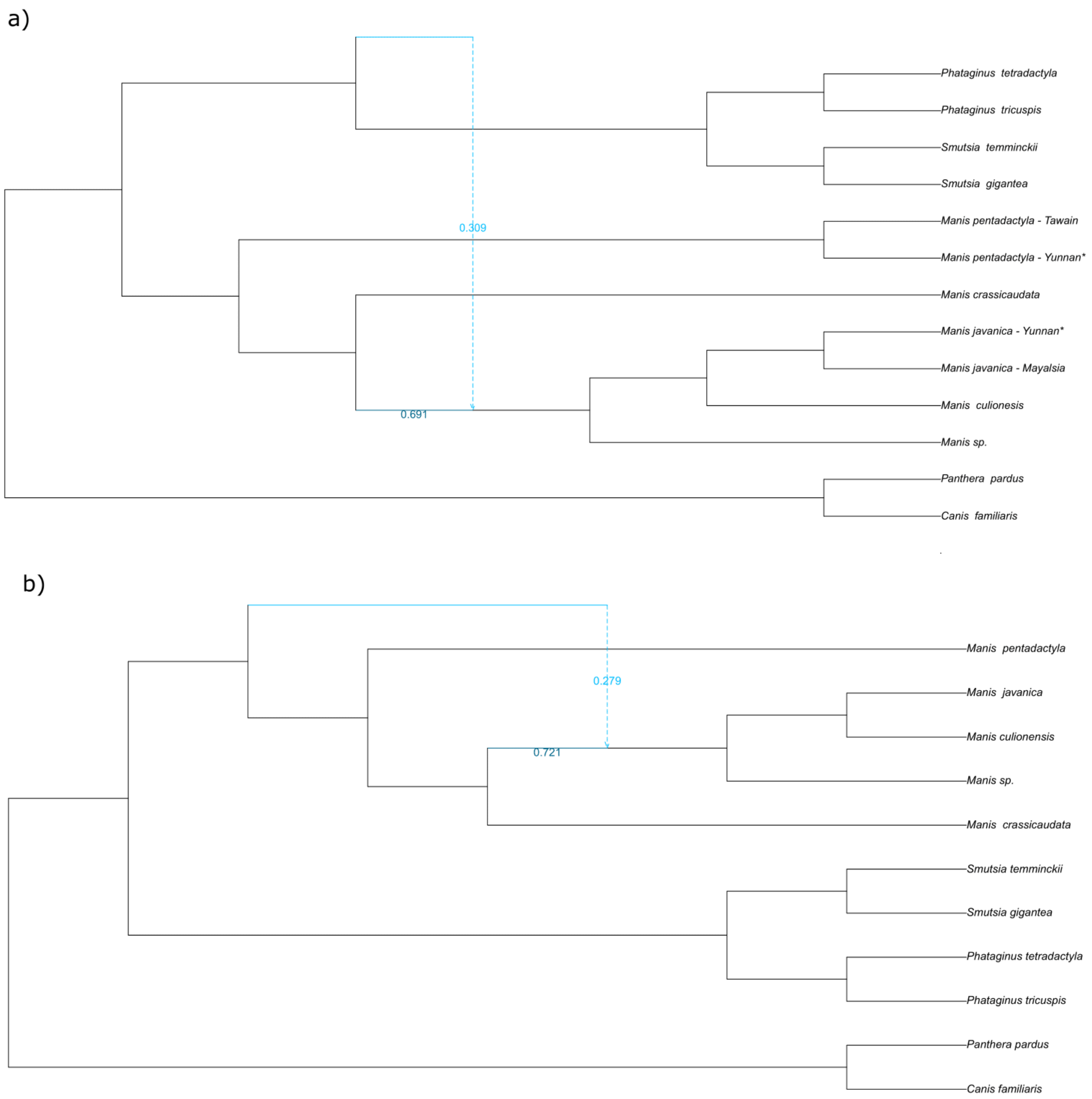


Figure S4: Maximum pseudolikelihood phylogentic networks

Phylogentic networks, using maximum pseudolikelihood estimates, depicted as rooted phylogenies to infer reticulation (introgression/hybridization) events within pangolins. Dotted blue lines indicate connection and direction of gene-flow between the donner and recipient taxa. Solid blue lines indicate ancestry of donner and recipient taxa. Numbers correspond to the proportion of genes shared between recipient and donner (light blue) and recipient and ancestor (dark blue). Analyses were run twice, (a) first by using each individual as a spate evolutionary unit and then (b) by indicating that multiple individuals of the same species were the same species. Both analyses indicated 1 reticulation event as the most likely outcome. Networks were drawn using the PhyloPlot function in Julia. Asterisks (*) indicate confiscated individuals whose origins could not be verified.

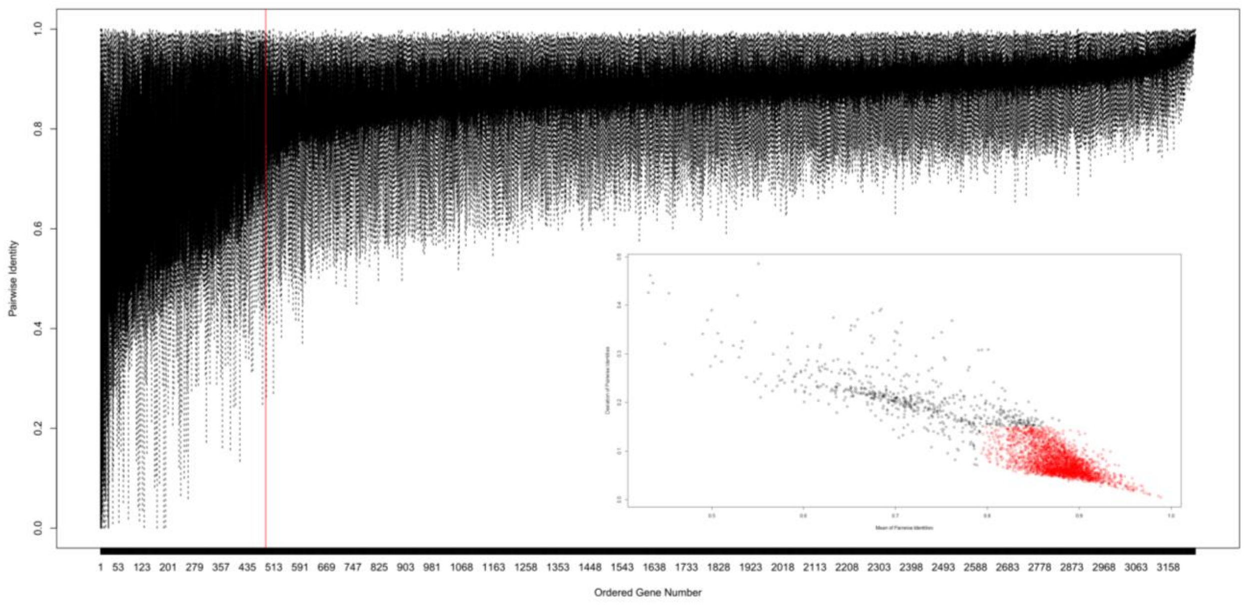


Figure S5: Diversity of orthologous whole-gene markers

Boxplot of whole-genes ranked from lowest (most diverse) to highest (least diverse) mean pairwise identity and the deviation of these markers (inner scatter plot). The average pairwise identity was calculated from the multiple sequence alignments of the eight pangolin species (using a single representative of each species only), by obtaining the pairwise identity for each combination of pairs and averaging it. This indicates the level of similarity (1=100% mean pairwise identity/similarity) which can be interpreted as pairwise diversity on the inverse (1=0% diverse). The vertical red line on the boxplot indicates the point (marker 566) at which genes to the left of it are likely outliers (values >2 Qn deviations from the median of the mean pairwise identities) as they are too diverse to be reliable (possible influence of parology, repetitive regions, bad alignment, bad gene annotation, etc.). This cutoff, along with another (values >2 Qn deviations from the median of the standard deviations of mean pairwise identities) can be viewed in the scatter plot whereby black points are the outliers (first 610 markers with the lowest mean pairwise identity or highest standard deviation of mean pairwise identity) and red points are likely reliable markers. These outlier markers have been highlighted in red in the sheet "removed genes" the diversity database (Database S1: available at https://doi.org/10.5281/zenodo.7517409). The rest make up 3 410 610 polymorphisms from 2 623 orthologous whole-genes.

**Table S1:** The samples sequenced and references used in this study. BUSCO v5 scores were obtained from mammalian orthologues by uploading the assemblies to gVolante (https://gvolante.riken.jp/). Unpublished genome assemblies and sequencing data for DNA Zoo versions of *Manis pentadactyla*, *M. javanica*, and *Phataginus tricuspis* are used with permission from the DNA Zoo Consortium (dnazoo.org). These DNA Zoo draft assemblies were created and reviewed following the Hi-C method (Dudchenko, et al. 2017; Dudchenko, et al. 2018).

| **Individual** | **Genbank accession number** | **Isolate; origin** | **Study** | **Collection year; sample type; collector** | **Sequencing platform** | **BUSCO scores (completed_S; completed_D; Fragmented; Missing)** | **Estimated coverage. For samples in this study: (1) using Lander/Waterman equation with 2.45 Gb genome length as standard / (2) mapped to *P. tricuspis* DNA Zoo reference.** | **Notes** |
|---|---|---|---|---|---|---|---|---|
| Chinese pangolin (*Manis pentadactyla*) | GCA_000738955.1 | MPE899; Taiwan | (Choo, et al. 2016) | N/A | Illumina HiSeq | 70.5; 0.7; 10; 18.8 | 59x | Unpublished DNA Zoo version with additional Hi-C and NovaSeq data can be found here: https://www.dnazoo.org/assemblies/Manis_pentadactyla |
| Chinese pangolin (*Manis pentadactyla*) | GCA_014570555.1 | MP20; Confiscated in Yunnan, China | (Hu, Hao, et al. 2020) | N/A | Illumina HiSeq and 10X genomic | 93.1; 1.3; 1.2; 4.4 | 281.6x | Used 77.19 Gb of the 180.66 Gb available data for mapping |
| Sunda pangolin (*Manis javanica*) | GCA_001685135.1 | MP_PG03-UM; Malaysia | (Choo, et al. 2016) | N/A | Illumina HiSeq | 74.6; 0.6; 6; 8.8 | 145.7x | Used 71.33 Gb of the 169.4 Gb available data for mapping Unpublished DNA Zoo version |

| | | | | | | | | |
|---|---|---|---|---|---|---|---|---|
| | | | | | | | | with additional Hi-C and HiSeq data can be found here: https://www.dnazoo.org/assemblies/Manis_javanica |
| Sunda pangolin (*Manis javanica*) | GCA_014570535.1 | MJ74; Confiscated in Yunnan, China | (Hu, Hao, et al. 2020) | N/A | Illumina HiSeq and 10X genomic | 92.3; 0.9; 2; 4.8 | 411.8x | Used 112.07 Gb of the 316.2 Gb available data for mapping |
| Philippine pangolin (*Manis culionensis*) | | MCUP0005; Casuyan, Palawan Isl., Philippines | This study | 1945; museum skin; H.H. Hoogstraal / Field Museum of Natural History, Chicago (FMNH 62919) | Illumina HiSeq X Ten | 6.2; 0; 12.5; 81.3 | (1) 76.7x / (2) 19.9x | |
| Indian pangolin (3) (*Manis crassicaudata*) | | MCR3; Confiscated but died in captivity, Sri Lanka | This study | 2008; tissue from dead individual; Jayanthi Alahakoon / Colombo Zoo | Illumina HiSeq X Ten | 40.9; 0.2; 16.9; 42 | (1) 53.1x / (2) 33.2x | |
| Indian pangolin (4) (*Manis crassicaudata*) | | MCR4; Confiscated but died in captivity, Sri Lanka | This study | 2008; tissue from dead individual; Jayanthi Alahakoon / Colombo Zoo | Illumina HiSeq X Ten | N/A | (1) 16.2x / (2) 9.1x | Only used in Heterozygosity and PSMC analyses due to limited coverage |
| *Manis sp.* | GCA_016801295.1 | Confiscated in Sichuan, China | (Cao, et al. 2021) | N/A | Illumina HiSeq | 54.5; 0.3; 13.9; 31.3 | 44x | Indicated at *M. crassicaudata* on NCBI |
| Black-bellied pangolin (*Phataginus tetradactyla*) | | CAM085; Yaoundé bushmeat market, Cameroon | This study | 2007; tissue from dead individual; F. | Illumina HiSeq 3000 | 55.9; 0.4; 15.3; 28.4 | (1) 43.4x / (2) 30.1x | |

| | | | | | | | | |
|---|---|---|---|---|---|---|---|---|
| | | | | Njiokou & P. Gaubert | | | | |
| White-bellied pangolin (*Phataginus tricuspis*) | GCA_004765945.1 | BS60 | Unpublished - BROAD institute | O. Ryder / San Diego Zoo Institute for Conservation Research | Illumina HiSeq | 65.5; 0.8; 12.4; 21.3 | 30.2x | |
| White-bellied pangolin (*Phataginus tricuspis*) | | Jaziri | Unpublished - DNA Zoo | Pittsburgh Zoo & PPG Aquarium | Illumina NovaSeq and Hi-C | 87.4; 1.5; 4.2; 6.9 | Unknown | The assembly can be found here courtesy of DNA Zoo: https://www.dnazoo.org/assemblies/Phataginus_tricuspis |
| Temminck's pangolin (*Smutsia temminckii*) | | STEM 81; Kalahari Oryx Game Farm, South Africa | This study | 2012; tissue from dead individual; D.W. Pietersen | Illumina HiSeq X Ten | 46.4; 0.4; 17.2; 36 | (1) 43.9x / (2) 21.9x | |
| Giant pangolin (*Smutsia gigantea*) | | CAM011; Yaoundé bushmeat market, Cameroon | This study | 2007; tissue from dead individual; F. Njiokou & P. Gaubert | Illumina HiSeq 3000 and Oxford Nanopore | 79.8; 1.3; 7; 11.9 | (1) 100.4x / (2) 57.4x | Mapping coverage estimate is from short read data only |

**Table S2**: Output of the concordance analysis implemented in IQ-TREE along with the significance of the Chi-squared test of independence between the two discordant gene counts (gDF1_N and gDF2_N). This was used to test whether incomplete lineage sorting (ILS) may be the sole cause of discordance for a branch whereby a significant p-value (*) indicates the contrary. Branch Clade refers to the clade onto which the branch opens in the phylogenetic tree in Figure 1. The rest of the column ID's follow that of IQ-TREE; ID: Branch ID, gCF: Gene concordance factor (=gCF_N/gN %), gCF_N: Number of trees concordant with the branch, gDF1: Gene discordance factor for NNI-1 branch (=gDF1_N/gN %), gDF1_N: Number of trees concordant with NNI-1 branch, gDF2: Gene discordance factor for NNI-2 branch (=gDF2_N/gN %), gDF2_N: Number of trees concordant with NNI-2 branch, gDFP: Gene discordance factor due to polyphyly (=gDFP_N/gN %), gDFP_N: Number of trees decisive but discordant due to polyphyly, gN: Number of trees decisive for the branch, sCF: Site concordance factor averaged over 100 quartets (=sCF_N/sN %), sCF_N: sCF in absolute number of sites, sDF1: Site discordance factor for alternative quartet 1 (=sDF1_N/sN %), sDF1_N: sDF1 in absolute number of sites, sDF2: Site discordance factor for alternative quartet 2 (=sDF2_N/sN %), sDF2_N: sDF2 in absolute number of sites, sN: Number of informative sites averaged over 100 quartet, gEF_p: p-value of the Chi-squared test of independence for genes.

| **Branch ID** | **Branch Clade** | **gCF** | **gCF_N** | **gDF1** | **gDF1_N** | **gDF2** | **gDF2_N** | **gDFP** | **gDFP_N** | **gN** | **sCF** | **sCF_N** | **sDF1** | **sDF1_N** | **sDF2** | **sDF2_N** | **sN** | **Branch-length** | **gEF_p** |
|---|---|---|---|---|---|---|---|---|---|---|---|---|---|---|---|---|---|---|---|
| 16 | *M. culionensis*/*M. javanica* Malaysia/*M. javanica* China | 91.32 | 2957 | 3.18 | 103 | 3.68 | 119 | 1.82 | 59 | 3238 | 86.62 | 77921.4 | 7.18 | 5998.11 | 6.21 | 5318.77 | 89238.28 | 2.13912 | 0.277353521 |
| 18 | *M. sp./M. culionensis/M. javanica Malaysia/M. javanica China* | 97.59 | 3160 | 0.12 | 4 | 0.06 | 2 | 2.22 | 72 | 3238 | 89.69 | 247931.1 | 5.21 | 12487.84 | 5.1 | 12177.43 | 272596.4 | 3.91262 | 0.419316578 |
| 19 | *M. crassicaudata/M. sp./M. culionensis/M. javanica* Malaysia/*M. javanica* China | 94.6 | 3063 | 2.5 | 81 | 1.14 | 37 | 1.76 | 57 | 3238 | 67.99 | 166950 | 16.91 | 39310.42 | 15.1 | 35605.53 | 241866 | 2.71664 | 4.98E-05* |
| 20 | *M. pentadactyla* Taiwan/*M. pentadactyla* China | 98.67 | 3195 | 0.37 | 12 | 0.37 | 12 | 0.59 | 19 | 3238 | 96.5 | 310196.3 | 1.59 | 4394.39 | 1.91 | 5194.61 | 319785.3 | 4.16025 | 1 |
| 21 | *Manis* (Asian pangolins) | 96.15 | 2373 | 1.22 | 30 | 1.22 | 30 | 1.42 | 35 | 2468 | 81.27 | 391309.6 | 8.93 | 42936.17 | 9.79 | 46976.18 | 481221.9 | 3.04534 | 1 |
| 22 | Pholidota & Carnivora | 93.71 | 1072 | 2.8 | 32 | 3.15 | 36 | 0.35 | 4 | 1144 | 91.01 | 1235125 | 4.27 | 57919.86 | 4.72 | 64050.99 | 1357096 | 2.38006 | 0.626943182 |

| | | | | | | | | | | | | | | | | | | | |
|---|---|---|---|---|---|---|---|---|---|---|---|---|---|---|---|---|---|---|---|
| 23 | African pangolins | 96.64 | 2385 | 1.01 | 25 | 1.26 | 31 | 1.09 | 27 | 2468 | 76.93 | 345520.9 | 11.82 | 53083.68 | 11.25 | 50536.63 | 449141.2 | 3.12795 | 0.409780382 |
| 24 | *Phataginus* | 96.54 | 3127 | 1.27 | 41 | 1.08 | 35 | 1.11 | 36 | 3239 | 79.05 | 248559.4 | 10.28 | 29904.15 | 10.67 | 31191.61 | 309655.2 | 3.13571 | 0.480161224 |
| 25 | *Smutsia* | 97.44 | 3155 | 0.65 | 21 | 0.8 | 26 | 1.11 | 36 | 3238 | 84.25 | 280467 | 8.01 | 24451.21 | 7.74 | 23795.52 | 328713.7 | 3.56152 | 0.479055921 |
| 17 | *M. javanica* Malaysia/*M. javanica* China | 90.15 | 2919 | 3.74 | 121 | 4.39 | 142 | 1.73 | 56 | 3238 | 88.39 | 23207.07 | 5.95 | 1502.69 | 5.66 | 1444.31 | 26154.07 | 2.02097 | 0.19449974 |

**Table S3:** Dates used for soft bound fossil calibrations on specific nodes to be used as priors for the MCMCtree analysis of divergence estimates of pangolins. These calibrations are based on both dated fossils and molecular phylogeny estimates with reasoning provided for each. The calibration of Pholidota is the most recent calibration node possible due to the scarcity of fossils for genus/species-based estimates of the group.

| **Node** | **Date** | **Fossil** | **Reference** | **Notes** |
|---|---|---|---|---|
| Ferae | 66–87 Ma | Min<br>UALVP 50993 and 50994 (Oldest stem-carnivores - miacids, viverravids)<br><br>Max<br>Molecular estimate | (Fox, et al. 2010)<br><br>(Zhou, et al. 2011;<br>Gaubert, et al. 2018) | Due to no upper estimates of Ferae, we used a molecular dated calibration which has been used in previous studied |
| Carnivora | 37.3–66 Ma | Min<br>*Daphoenus* & *Hesperocyon*<br><br>Max<br>UALVP 50993 and 50994 (Oldest stem-carnivores - miacids, viverravids) | (Benton, et al. 2015)<br><br>(Fox, et al. 2010) | *Tapocyon* may be an even older caniform; (46–43 Ma). However, it is placed outside Carnivora (Wesley-Hunt and Flynn 2005). The oldest feliforms may be the nimravids, but this too is uncertain (Hunt 2004). |
| Pholidota | 31–45 Ma | Min<br>*Manidae* (oldest manidae fossil)<br><br>Max<br>*Euromanis krebsi* (oldest)<br>*Eurotamandua joresi*<br>*Eomanis waldi* | (Gebo and Rasmussen 1985)<br><br>(Gaudin, et al. 2009; Rose 2012) | Messel deposits (Germany). Found with *Eurotamandua joresi* which has been debated as to whether it should be included in the Pholidota or whether it predates this order but Gaudin, et al. (2009) places it under Pholidota as sister to *Eomanis* and *Euromanis* |

**Table S4:** Estimated posterior mean or median divergence estimates and 95% Highest Posterior Density (HPD) interval of each node for this study and the one conducted by Gaubert, et al. (2018). The latter study involved mitogenomes and nine nuclear genes, included more individuals and fossil calibrations within the order Carnivora, and used the program BEAST (Bouckaert, et al. 2014) to obtain time to most recent common ancestor (TMRCA) estimates. $ refers to nodes with fossil priors for this study. * refers to nodes where the 95% HPD of divergence estimates do not overlap in the two studies.

| | **This study** | | **Gaubert et al. (2018)** | |
|---|---|---|---|---|
| **Node** | **Mean** | **95% HPD** | **Median** | **95% HPD** |
| Ferae$ | 79.47 | 67.66 – 87.24 | 78.9 | 69.6 – 87.0 |
| Carnivora$ | 49.95 | 36.49 – 65.44 | 50.8 | 44.9 – 57.4 |
| Caniformia | | | 41.6 | 38.0 – 46.0 |
| *Mustela – Ailuropoda* | | | 35.4 | 30.5 – 40.9 |
| Felidae | | | 11.3 | 10.0 – 13.3 |
| *Acinonyx – Felis* | | | 7.2 | 3.3 – 10.2 |
| Pholidota$ | 41.34 | 33.54 – 45.45 | 37.9 | 31.4 – 44.6 |
| *Smutsia - Phataginus* | 20.35 | 12.79 – 28.29 | 22.9 | 18.7 – 27.2 |
| *Manis* | 22.62 | 17.07 – 28.23* | 12.9 | 10.3 – 15.6 |
| *M. crassicaudata – (M. sp, M. javanica, M. culionensis)* | 16.84 | 12.21 – 22.00* | 9.1 | 6.6 – 11.4 |
| *M. sp – (M. javanica – M. culionensis)* | 7.16 | 4.73 – 10.42 | | |
| *M. javanica – M. culionensis* | 2.71 | 1.70 – 4.21 | 1.7 | 0.4 – 2.5 |
| *M. javanica* China – *M. javanica* Malaysia | 0.77 | 0.46 – 1.24 | | |
| *M. pentadactyla* China – *M. pentadactyla* Taiwan | 2.82 | 1.48 – 4.95 | | |
| *Smutsia* | 9.78 | 5.55 – 15.74 | 9.8 | 5.6 – 13.2 |
| *Phataginus* | 11.34 | 6.55 – 17.45 | 13.3 | 9.3 – 16.5 |
| *P. tricuspis* | | | 2.7 | 0.8 – 4.6 |
| *P. tricuspis* West Africa – Western Central Africa | | | 1.1 | 0.0 – 2.4 |

**Table S5:** Proportion of genome-wide heterozygosity for mammalian species ranked from least to most diverse. As depicted in Figure 4, each mammalian order is colour-coded (Pholidota = dark red), and citations are given for the studies in which the value was originally provided for each species. Heterozygosity values for pangolins (Pholidota) include those calculated in our study as well as Hu, Hao, et al. (2020), who used a different method of calculation (using VCFtools on autosomal SNPs with 50 kb non-overlapping sliding windows). This is an edited and updated version of the table created by Hu, Hao, et al. (2020).

| **Species** | **Heterozygosity (%)** | **Mammalian order** | **Sources** |
|---|---|---|---|
| Iberian lynx (*Lynx pardinus*) | 0.010 | Carnivora | (Abascal, et al. 2016) |
| Domestic cat (*Felis catus*) | 0.012 | Carnivora | (Cho, et al. 2013) |
| Baiji (*Lipotes vexillifer*) | 0.012 | Certartiodactyla | (Zhou, et al. 2013) |
| Myanmar snub-nosed monkey (*Rhinopithecus strykeri*) | 0.015 | Primates | (Zhang, et al. 2016) |
| Altai Neanderthal (*Homo sapiens*) | 0.017 | Primates | (Prüfer, et al. 2014) |
| Cheetah (*Acinonyx jubatus*) | 0.020 | Carnivora | (Dobrynin, et al. 2015) |
| Snow leopard (*Panthera uncia syn*) | 0.023 | Carnivora | (Cho, et al. 2013) |
| Yangtze river dolphin (*Lipotes vexillifer*) | 0.026 | Certartiodactyla | (Zhou, et al. 2013) |
| Eurasian lynx (*Lynx lynx*) | 0.028 | Carnivora | (Abascal, et al. 2016) |
| Siberian tiger (*Panthera tigris altaica)* | 0.030 | Carnivora | (Dobrynin, et al. 2015) |
| Domestic dog (*Canis familiaris*) | 0.032 | Carnivora | (Lindblad-Toh, et al. 2005) |
| Tasmanian devil (*Sarcophilus harrisii*) | 0.032 | Marsupialia | (Cho, et al. 2013) |
| Black snub-nosed monkey (*Rhinopithecus bieti*) | 0.033 | Primates | (Zhou, et al. 2016) |
| Island fox (*Urocyon littoralis*) - San Miguel | 0.033 | Carnivora | (Robinson, et al. 2016) |
| Bengal tiger (*Panthera tigris tigris*) | 0.040 | Carnivora | (Dobrynin, et al. 2015) |
| Golden snub-nosed monkey (*Rhinopithecus roxellana*) | 0.042 | Primates | (Zhou, et al. 2016) |
| White lion (*Panthera leo*) | 0.048 | Carnivora | (Cho, et al. 2013) |
| Amur tiger (*Panthera tigris altaica*) | 0.049 | Carnivora | (Cho, et al. 2013) |
| Island fox (*Urocyon littoralis*) - San Nicolis | 0.049 | Carnivora | (Robinson, et al. 2016) |
| Aye-Aye (*Daubentonia madagascariensis*) | 0.051 | Primates | (Perry, et al. 2011) |
| Wild horse (*Equus ferus przewalskii*) | 0.052 | Perissodactyla | (Huang, et al. 2014) |
| African lion (*Panthera leo*) | 0.058 | Carnivora | (Cho, et al. 2013) |
| Grey snub-nosed monkey (*Rhinopithecus brelichi*) | 0.062 | Primates | (Zhou, et al. 2016) |
| Eastern lowland gorilla (*Gorilla beringei graueri*) | 0.064 | Primates | (Xue, et al. 2015) |
| Bornean orangutan (*Pongo pygmaeus*) | 0.065 | Primates | (Locke, et al. 2011) |
| Mountain gorilla (*Gorilla beringei beringei*) | 0.065 | Primates | (Xue, et al. 2015) |
| Naked mole rat (*Heterocephalus glaber*) | 0.068 | Rodentia | (Kim, et al. 2011) |

| | | | |
|---|---|---|---|
| Pileated gibbon (*Hylobates pileatus*) | 0.073 | Primates | (Carbone, et al. 2014) |
| White tiger (*Panthera tigris tigris*) | 0.073 | Carnivora | (Cho, et al. 2013) |
| White-bellied pangolin (Phataginus tricuspis) - DG | 0.074 | Pholidota | Sequencing data from DNAZoo |
| Dromedary (*Camelus dromedarius*) | 0.074 | Certartiodactyla | (Wu, et al. 2014) |
| Platypus (*Ornithorhynchus anatinus*) | 0.075 | Monotremata | (Warren, et al. 2008) |
| Indian pangolin (*Manis crassicaudata*) – MCR4 | 0.075 | Pholidota | This study |
| Eastern lowland gorilla (*Gorilla beringei graueri*) | 0.076 | Primates | (Scally, et al. 2012) |
| Human Han (*Homo* species) | 0.077 | Primates | (Meyer, et al. 2012) |
| West African chimpanzees (*Pan troglodytes verus*) | 0.080 | Primates | (Mikkelsen, et al. 2005) |
| African green monkey (*Chlorocebus aethiops aethiops*) | 0.080 | Primates | (Warren, et al. 2015) |
| Wild bactrian camel (*Camelus bactrianus ferus*) | 0.084 | Certartiodactyla | (Wang, et al. 2012) |
| Chinese pangolin (*Manis pentadactyla*) - Taiwan | 0.085 | Pholidota | Sequencing data from Choo, et al. (2016). Analysis from this study |
| Sunda pangolin (*Manis javanica*) - Average over 74 individuals | 0.085 | Pholidota | Samples in Hu, Hao, et al. (2020). Analysis in their study |
| Minke whale (*Balaenoptera acutorostrata*) | 0.086 | Certartiodactyla | (Yim, et al. 2014) |
| Finless porpoise (*Neophocaena phocaenoides*) | 0.086 | Certartiodactyla | (Yim, et al. 2014) |
| Koala (*Phascolarctos cinereus*) | 0.087 | Marsupialia | (Johnson, et al. 2018) |
| Tibetan antelope (*Pantholops hodgsonii*) | 0.088 | Certartiodactyla | (Ge, et al. 2013) |
| Yak (*Bos grunniens*) | 0.089 | Certartiodactyla | (Qiu, et al. 2012) |
| Mongolian horse (*Equus ferus caballus*) | 0.089 | Perissodactyla | (Huang, et al. 2014) |
| Domestic bactrian camel (*Camelus bactrianus*) | 0.090 | Certartiodactyla | (Wang, et al. 2012) |
| Southern white rhinoceros (*Ceratotherium simum simum*) | 0.090 | Perissodactyla | (Tunstall, et al. 2018) |
| Indian pangolin (*Manis crassicaudata*) – MCR3 | 0.094 | Pholidota | This study |
| Common chimpanzee (*Pan troglodytes*) | 0.095 | Primates | (Mikkelsen, et al. 2005) |
| Domestic horse (*Equus ferus caballus*) | 0.095 | Perissodactyla | (Wade, et al. 2009) |
| Cross River gorilla (*Gorilla gorilla diehli*) | 0.096 | Primates | (Xue, et al. 2015) |
| Black-bellied pangolin (*Phataginus tetradactyla*) | 0.100 | Pholidota | This study |
| Wrangel woolly mammoth (*Mammuthus primigenius*) | 0.100 | Proboscidea | (Palkopoulou, et al. 2015) |
| Polar bear (*Ursus maritimus*) | 0.108 | Carnivora | (Liu, et al. 2014) |
| Chinese pangolin (*Manis pentadactyla*) – Yunnan, China confiscation | 0.109 | Pholidota | Sequencing data from Hu, Hao, et al. (2020). |

| | | | |
|---|---|---|---|
| | | | Analysis from this study |
| Island fox (*Urocyon littoralis*)- San Clemente | 0.110 | Carnivora | (Robinson, et al. 2016) |
| Northern white rhinoceros (*Ceratotherium simum cottoni*) | 0.110 | Perissodactyla | (Tunstall, et al. 2018) |
| Chinese pangolin (*Manis pentadactyla*) | 0.114 | Pholidota | Sample MP20 in Hu, Hao, et al. (2020). Analysis in their study |
| Bactrian camel (*Camelus bactrianus*) | 0.116 | Certartiodactyla | (Wu, et al. 2014) |
| Sumatran orangutan (*Pongo abelii*) | 0.120 | Primates | (Xue, et al. 2015) |
| Gray fox (*Urocyon cinereoargenteus*) | 0.120 | Carnivora | (Robinson, et al. 2016) |
| Brown hyena (*Parahyaena brunnea*) | 0.121 | Carnivora | (Westbury, et al. 2018) |
| Cow (*Bos taurus*) | 0.121 | Certartiodactyla | (Corbett-Detig, et al. 2015) |
| Rat (*Rattus norvegicus*) | 0.125 | Rodentia | (Leffler, et al. 2012) |
| Oimyakon woolly mammoth (*Mammuthus primigenius*) | 0.125 | Proboscidea | (Palkopoulou, et al. 2015) |
| Chinese pangolin (*Manis pentadactyla*) - Average over 23 individuals | 0.127 | Pholidota | Samples in Hu, Hao, et al. (2020). Analysis in their study |
| Sumatran rhinoceros (*Dicerorhinus sumatrensis*) | 0.130 | Perissodactyla | (Mays, et al. 2018) |
| Giant panda (*Ailuropoda melanoleuca*) | 0.132 | Carnivora | (Li, et al. 2010) |
| Siamang (*Symphalangus syndactylus*) | 0.140 | Primates | (Carbone, et al. 2014) |
| Bottlenose dolphin (*Tursiops truncatus*) | 0.142 | Certartiodactyla | (Yim, et al. 2014) |
| Western lowland gorilla (*Gorilla gorilla gorilla*) | 0.144 | Primates | (Xue, et al. 2015) |
| Giant pangolin (*Smutsia gigantea*) | 0.146 | Pholidota | This study |
| Gray wolf (*Canis lupus*) | 0.149 | Carnivora | (Corbett-Detig, et al. 2015) |
| Fin whale (*Balaenoptera physalus*) | 0.151 | Certartiodactyla | (Yim, et al. 2014) |
| Sunda pangolin (*Manis javanica*) | 0.152 | Pholidota | Sample MJ74 in Hu, Hao, et al. (2020). Analysis in their study |
| Temminck's pangolin (*Smutsia temminckii*) | 0.155 | Pholidota | This study |
| Chinese hamster (*Cricetulus griseus*) | 0.159 | Rodentia | (Lewis, et al. 2013) |
| Asian pangolin sp. (Manis sp.) - Sichuan, China confiiscated | 0.161 | Pholidota | Sequencing data from Cao, et al. (2021). Analysis in this study. |
| Silvery gibbon (*Hylobates moloch*) | 0.17 | Primates | (Carbone, et al. 2014) |
| Malaysian cynomolgus macaque (*Macaca fascicularis*) | 0.171 | Primates | (Higashino, et al. 2012) |
| Central African chimpanzees (*Pan troglodytes troglodytes*) | 0.176 | Primates | (Mikkelsen, et al. 2005) |

| | | | |
|---|---|---|---|
| Western lowland gorilla (*Gorilla gorilla gorilla*) | 0.178 | Primates | (Scally, et al. 2012) |
| Vervet monkey (*Chlorocebus aethiops pygerythrus*) | 0.180 | Primates | (Warren, et al. 2015) |
| Wild boar (*Sus scrofa*) - Tibetan | 0.182 | Certartiodactyla | (Li, et al. 2013) |
| Olive baboon (*Papio anubis*) | 0.189 | Primates | (Corbett-Detig, et al. 2015) |
| Island fox (*Urocyon littoralis*) - San Rosa | 0.191 | Carnivora | (Robinson, et al. 2016) |
| Island fox (*Urocyon littoralis*) - San Catalina | 0.196 | Carnivora | (Robinson, et al. 2016) |
| Island fox (*Urocyon littoralis*) - San Cruz | 0.197 | Carnivora | (Robinson, et al. 2016) |
| Northern white-cheeked gibbon (*Nomascus leucogenys*) | 0.220 | Primates | (Carbone, et al. 2014) |
| Bighorn sheep (*Ovis canadensis*) | 0.222 | Certartiodactyla | (Corbett-Detig, et al. 2015) |
| Sunda pangolin (*Manis javanica*) - Malaysia | 0.224 | Pholidota | Sequencing data from Choo, et al. (2016). Analysis from this study |
| Sunda pangolin (*Manis javanica*) - Yunnan, China confiscation | 0.230 | Pholidota | Sequencing data from sample MJ74 in Hu, Hao, et al. (2020). Analysis from this study |
| Alpaca (*Vicugna pacos*) | 0.266 | Certartiodactyla | (Wu, et al. 2014) |
| David’s myotis (*Myotis davidii*) | 0.279 | Chiroptera | (Zhang, et al. 2013) |
| Rhesus macaque (*Macaca mulatta*) | 0.287 | Primates | (Corbett-Detig, et al. 2015) |
| Cape buffalo (*Syncerus caffer caffer*) | 0.288 | Certartiodactyla | (de Jager, et al. 2021) |
| Brown bear (*Ursus arctos*) | 0.320 | Carnivora | (Liu, et al. 2014) |
| White-bellied pangolin (*Phataginus tricuspis*) - CWA | 0.334 | Pholidota | Sequencing data from Genbank (GCA_004765945.1). Analysis from this study |
| Common marmoset (*Callithrix jacchus*) | 0.341 | Primates | (Consortium 2014) |
| Przewalski's horse (*Equus ferus przewalskii*) | 0.363 | Perissodactyla | (Corbett-Detig, et al. 2015) |
| Brandt’s bat (*Myotis brandtii*) | 0.371 | Chiroptera | (Seim, et al. 2013) |
| Chinese rhesus macaque (*Macaca mulatta lasiota*) | 0.410 | Primates | (Yan, et al. 2011) |
| Wild boar (*Sus scrofa*) | 0.441 | Certartiodactyla | (Corbett-Detig, et al. 2015) |
| Black flying fox (*Pteropus alecto*) | 0.453 | Carnivora | (Zhang, et al. 2013) |
| Opossum (*Monodelphis domestica*) | 0.490 | Marsupialia | (Mikkelsen, et al. 2007) |
| Philippine pangolin (*Manis culionensis*) | 0.492 | Pholidota | This study |

| | | | |
|---|---|---|---|
| Crab-eating macaque (*Macaca fascicularis*) | 0.530 | Primates | (Yan, et al. 2011) |
| Rabbit (*Oryctolagus cuniculus*) | 0.750 | Lagomorpha | (Carneiro, et al. 2014) |
| Eastern hoolock gibbon (*Hoolock leuconedys*) | 0.800 | Primates | (Carbone, et al. 2014) |
| House mouse (*Mus musculus castaneus*) | 0.809 | Rodentia | (Corbett-Detig, et al. 2015) |

**Table S6:** Estimated generation times of each species of pangolins based on the currently available literature. These times include a sum of the gestation period (passing on genomic information to the next generation at conception) and time until sexual maturity (when genomic information can be passed on to the next generation again). These estimates were used in the PSMC (pairwise sequentially Markovian coalescent) model analysis on pangolins in order to get the timing of changes in IICR (inverse instantaneous coalescence rate) as accurate as possible. We defined generation time as from the conception of an individual to the time until the first born of that individual is conceived.

| Species | Estimated generation time | Evidence | Citations |
|---|---|---|---|
| Chinese pangolin (*Manis pentadactyla*) | 2 years | Reaches sexual maturity at 1–1.5 years old, but may be as a low as 6–7 months; gestation is 6–7 months (captive data) | (Chin, et al. 2012; Zhang, et al. 2016) |
| Sunda pangolin (*Manis javanica*) | 1.5 years | 1 year (supposed sexual maturity of males is 1.5 years based off sperm analysis – A. Kurniawan, unpublished) until sexual maturity but may be as low as 6–7 months; gestation is 6 months (captive and seizure data) | (Zhang, et al. 2015; Zhang, et al. 2017) |
| Philippine pangolin (*Manis culionensis*) | 1.5 years | No data, but likely similar to the Sunda pangolin. | |
| Indian pangolin (*Manis crassicaudata*) | 3 years | Up to 3 years until sexual maturity (unpublished data), gestation is around 6–8 months (135–251 days; captive data). | (Mohapatra, et al. 2018; Mahmood, et al. 2020) |
| Black-bellied pangolin (*Phataginus tetradactyla*) | 2.5 years | 2 years (unpublished suggestion – book chapter), gestation is 5 months (150 days; wild data). | (Pagès 1972a; Gudehus, et al. 2020) |
| White-bellied pangolin (*Phataginus tricuspis*) | 2.5 years | No data for maturity (probably similar to Black-bellied pangolin), gestation is 5 months (140–150 days; wild data). | (Pagès 1972b) |
| Temminck's pangolin (*Smutsia temminckii*) | 2.5 years | Sexual maturity is likely around 2 years but no home range is established for a few more years (assuming home range is important, then 3–7 years), gestation is 3.5–4.5 months (105–140 days; both captive and wild data). | (van Ee 1966; Pietersen, et al. 2020) |
| Giant pangolin (*Smutsia gigantea*) | 3 years | No data available, but likely similar, if not longer than, Temmincks's pangolin due to larger size. | |
